# A nuclear trafficking checkpoint licenses resolution of RNA–DNA hybrids at stalled RNAPII to maintain genome stability under genotoxic stress

**DOI:** 10.1101/2025.08.13.670051

**Authors:** Jérémy Sandoz, Clara Capelli, Philippe Catez, Anthony Hannus, Alice Brion, Lise-Marie Donnio, Pierre-Olivier Mari, Jean-Paul Concordet, Elisa Bergamin, Emmanuel Compe, Jean-Marc Egly, Frédéric Coin

**Affiliations:** Institut de Génétique et de Biologie Moléculaire et Cellulaire, Illkirch Cedex, C.U. Strasbourg, France; Centre National de la Recherche Scientifique, UMR7104, Illkirch, France; Institut National de la Santé et de la Recherche Médicale, U1258, Illkirch, France; Université de Strasbourg, Strasbourg, France; Laboratoire Structure et Instabilité des Génomes, INSERM U1154, CNRS UMR7196, Muséum national d’Histoire naturelle, 43 rue Cuvier, 75005, Paris, France; Institut NeuroMyoGène-Laboratoire Physiopathologie et Génétique du Neurone et du Muscle (INMG_PGNM), CNRS UMR 5261, INSERM U1315, Université de Lyon, Université Claude Bernard Lyon 1, Lyon, France

**Keywords:** nuclear protein import, DNA Damage Response, EXD2, R-loop

## Abstract

Genotoxic stress inhibits transcription, generating RNA–DNA hybrid intermediates that promote toxic transcription–replication conflicts (TRCs). Here, we show that genotoxic stress triggers transient Importin-α/β1-dependent nuclear import of the mitochondrial exonuclease EXD2 to resolve these conflicts and maintain genome stability. Mechanistically, once imported into the nucleus, EXD2 is recruited to elongation-arrested RNA polymerase II (RNAPII) associated with RNA–DNA hybrids, and its exonuclease activity limits their accumulation, thereby suppressing TRCs. Mutation of a putative EXD2 nuclear localization signal phenocopies disruption of nuclear import, whereas forced nuclear targeting of EXD2 bypasses it. However, sustained nuclear localization of EXD2 leads to mitotic defects, underscoring the importance of its tight spatial regulation. Consistent with a direct role for EXD2 in RNA–DNA hybrid resolution, overexpression of the ribonuclease RNaseH1 compensates for its depletion following DNA damage. Together, our findings introduce the concept of a trafficking-dependent checkpoint that transiently licenses EXD2 nuclease activity in the nucleus in response to genotoxic stress to resolve RNA–DNA hybrid intermediates at stalled RNAPII complexes

**Teaser:** Stress-triggered nuclear import of EXD2 resolves harmful RNA–DNA hybrids to protect genome stability after DNA damage.

## Introduction

In response to genotoxic stress, cells activate the DNA damage response (DDR), a coordinated signaling network that detects DNA lesions, halts cell cycle progression, and promotes DNA repair to preserve genome integrity (*1*), (*2*), (*3*), (*4*). Distinct repair pathways are mobilized depending on the nature of the lesion and the cell cycle stage. Among these, nucleotide excision repair (NER) removes bulky DNA adducts such as UV-induced pyrimidine (6–4) photoproducts (6–4PPs) (*5*), (*6*), (*7*). NER operates through two sub-pathways: global genome NER (GG-NER), which surveys the entire genome, and transcription-coupled NER (TC-NER), which selectively removes lesions that block elongating RNAPII on actively transcribed genes (*8*), (*9*), (*10*), (*11*).

Concomitant with DNA repair activation, cells rapidly repress RNAPII-dependent transcription genome-wide (*12*), (*13*), (*14*). This transcriptional shutdown persists until DNA lesions are removed, after which transcription is progressively restored through a regulated process termed recovery of RNA synthesis (RRS) (*9*). While transcriptional repression after DNA damage has been extensively characterized, the mechanisms that actively restore productive transcription remain incompletely understood (*15*), (*16*), (*17*), (*18*), (*19*). Increasing evidence indicates that transcription restart requires the removal of R-loop–like structures, operationally defined as RNaseH1-sensitive RNA–DNA hybrid intermediates, associated with stalled RNAPII complexes. These structures can interfere with DNA replication, generate toxic transcription–replication conflicts (TRCs), and promote genome instability if not efficiently resolved (*20*), (*21*). Failure to properly clear such RNA–DNA hybrids has been linked to defective transcription recovery in several DNA repair–deficient contexts, suggesting that processing of these structures represents a critical regulatory step in the transition from transcriptional shutdown to transcription restart (*22*), (*23*).

Although most DDR factors are constitutively nuclear, several proteins undergo regulated nucleocytoplasmic relocalization in response to genotoxic stress (*24*), (*25*), (*26*), (*27*). Nuclear protein import, primarily mediated by the Importin-α/β1 pathway, provides a mechanism to dynamically shuttle proteins to the nucleus under stress conditions, specifically when their constitutive nuclear presence could be detrimental. While stress-induced nuclear import has been reported for certain DNA repair factors, its functional contribution to the DDR remains poorly defined (*28*). More specifically, whether regulated nuclear trafficking acts as a checkpoint coordinating transcription restart with genome maintenance has not been directly addressed.

The RNA/DNA exonuclease EXD2 represents a candidate effector of such a mechanism. Initially characterized as a mitochondrial nuclease involved in mitochondrial genome maintenance (*29*), (*30*), EXD2 has more recently been implicated in nuclear processes, including homologous recombination repair (HRR) and recovery of transcription following genotoxic stress (*31*), (*32*), (*17*). However, whether EXD2 dynamically accesses the nucleus in response to DNA damage and how this relocalization contributes to transcription restart remain unresolved.

Here, we demonstrate that genotoxic stress triggers transient Importin-α/β1–dependent nuclear import of EXD2, enabling its recruitment to elongation-arrested RNAPII complexes. We show that nuclear availability of EXD2 is required for cell survival following DNA damage and that enforced nuclear targeting of nuclease-active EXD2 bypasses the requirement for stress-induced import. Mechanistically, our data indicate that nuclear EXD2 promotes resolution of transcription-associated R-loop-like RNA–DNA hybrids that accumulate during transcriptional shutdown, thereby facilitating transcription restart and limiting TRCs. However, sustained nuclear localization of EXD2 leads to mitotic defects, underscoring the importance of its spatial regulation.

We observed that EXD2 import is a broad mechanism triggered by transcription-blocking DNA lesions such as those generated by UV irradiation or by the chemotherapeutic agent cisplatin. Together, these findings uncover a new spatial licensing mechanism in the DNA damage response in which stress-triggered nuclear trafficking preserves genome stability by promoting the resolution of RNA–DNA hybrids.

## Results

### Stress-induced nuclear import is required for transcription recovery

To assess whether Importin-α/β1–dependent nuclear import plays a role in the cellular response to genotoxic stress, we first examined the UV sensitivity of U-2 OS cells treated with IMP, INI-43, and IVM, three mechanistically distinct nuclear import inhibitors targeting Importin-β1 (IMP and INI-43) or its interaction with Importin-α (IVM) (*33*), (*34*), (*35*). Each treatment significantly reduced cell survival after UV irradiation compared with DMSO-treated controls (Fig. 1a and Sup. Fig. 1a-b). The extent of UV hypersensitivity was comparable to that of cells depleted for the TC-NER factor CSB, but less severe than that observed upon depletion of the GG-NER factor XPC (Sup. Fig. 1b), suggesting that impaired nuclear import phenocopies defects in TC-NER rather than GG-NER.

**Fig. 1.**
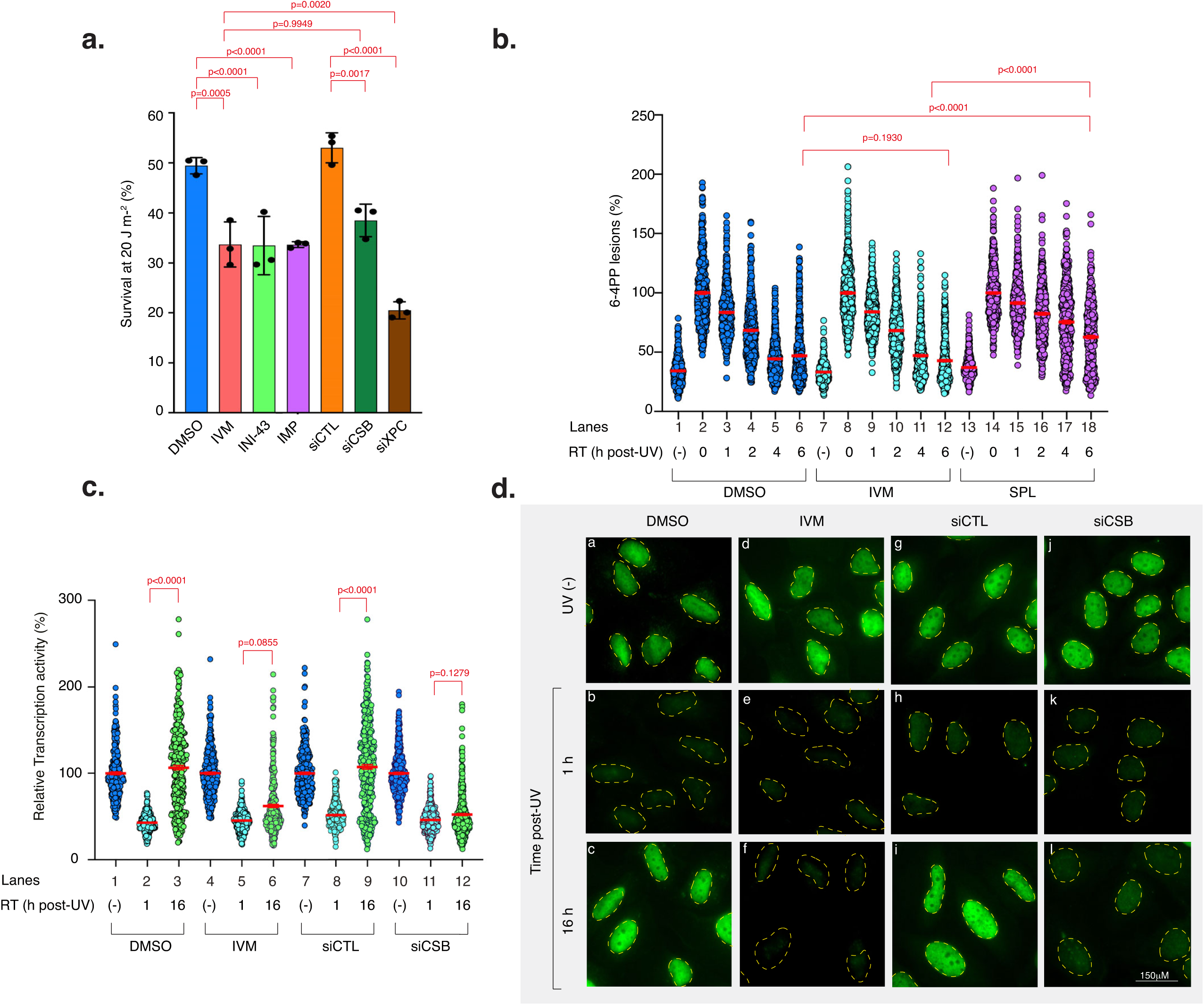
Nuclear import inhibition impairs transcription recovery but preserves DNA repair (a), Cell survival of U-2 OS cells treated with UV irradiation (20 J m^-2^). When indicated cells were treated for 1 h with inhibitors of nuclear import (1 μM) or with siRNA against the TC-NER factor CSB or against the GG-NER factor XPC for 48 h before UV irradiation. Cells were allowed to recover for 48 h and survival was determined. Data were normalized to the non-irradiated controls (set to 100%). Data are mean ± s.e.m from three independent experiments**. (b),** Removal of UV lesions was measured in U-2 OS cells, harvested at different time points post-UV (20 J m^-2^). Before UV irradiation cells were treated for 1 h with IVM (1μM) or for 2 h with SP (10 μM). Cells were labeled with anti-6-4PP antibody and signals were quantified using ImageJ at the different time points indicated in the figure. Graph represents the % of lesions remaining in the genome at different time points normalized to the amounts of lesions measured immediately after UV irradiation (as a value of 100%). Red bars indicate mean integrated densities ± s.e.m (*n*> 300 cells per condition from three independent experiments). RT; recovery time. (-); cells were mock-irradiated. 0; cells were UV-irradiated and fixed immediately. **(c),** Transcription rate in U-2 OS cells mock-or UV-irradiated at 20 J m^-2^. mRNA was labeled with EU at the indicated time points post-UV. When indicated, cells were treated for 1 h with IVM (1 μM) or for 48 h with siCTL or siCSB, before UV irradiation. To ensure specificity for RNAPII, cells were pre-treated with a low dose of actinomycin D to inhibit RNAPI transcription. EU signal was quantified by Fiji (ImageJ) and relative integrated densities, normalized to mock-treated level set to 100%, are reported on the graph. Red bars indicate mean integrated densities ± s.e.m (*n*> 300 cells per condition from three independent experiments). RT; recovery time. (-); cells were mock-irradiated. **(d),** Representative microscopy images of U-2 OS cells treated as in **(c)**. Statistical analyses were performed using one-way ANOVA with Tukey’s post hoc test.

We next addressed whether import inhibition affects the removal of UV-induced DNA lesions. Quantification of 6-4PPs (*16*), (*36*) revealed that IVM-treated cells removed lesions with kinetics indistinguishable from control cells (Fig. 1b). In contrast, the NER inhibitor spironolactone (*36*) markedly impaired lesion removal, validating the sensitivity of the assay. These results indicate that import inhibition does not compromise the excision repair of UV photoproducts, and therefore that the enhanced UV sensitivity is unlikely to stem from defective DNA repair.

Because the UV sensitivity profile of import-inhibited cells resembled that of CSB-deficient cells, we next examined whether nuclear import was required for transcription recovery following UV-irradiation. Using 5-ethynyluridine (EU) incorporation, we monitored nascent mRNA synthesis at different time points after UV irradiation. In both DMSO-and IVM-treated cells, global transcription was strongly repressed during the early hours following UV exposure (Fig. 1c-d). However, while transcription fully recovered by 16 h in control cells, it remained severely impaired in IVM-treated cells and comparable to that observed in CSB-depleted cells. Our experiments also indicated that transcription was not recovered in IVM-treated cells even 24 h after UV exposure, indicating a failure rather than a delay of transcription recovery (Sup. Fig. 1c).

These findings show that Importin-α/β1–dependent import is dispensable for DNA lesion recognition and removal, yet crucial for the reactivation of RNAPII transcription following genotoxic stress.

### Genotoxic stress induces Importin-α/β1–dependent nuclear import of EXD2

EXD2, a 3′–5′ exonuclease containing an N-terminal mitochondrial targeting sequence (MTS) followed by the catalytic exonuclease domain (ED) (Fig. 2a), has been the subject of considerable controversy regarding its subcellular localization and function. Initially characterized as a mitochondrial nuclease involved in mtRNA processing and mitochondrial genome maintenance, EXD2 was subsequently proposed to operate in the nucleus, where it contributes to homologous recombination repair and, more recently, to the recovery of transcription following UV irradiation (*30*), (*37*), (*31*), (*17*). These conflicting observations have left unresolved whether EXD2 acts exclusively as a mitochondrial nuclease or whether it participates more broadly in nuclear stress responses. In particular, no consensus exists as to whether EXD2 is capable of dynamic nucleocytoplasmic shuttling or whether it can access the transcription machinery in a damage-dependent manner. Given our findings above showing that transcription recovery requires Importin-α/β1–dependent nuclear import, the possible involvement of EXD2 in this process warranted further investigation.

**Fig. 2.**
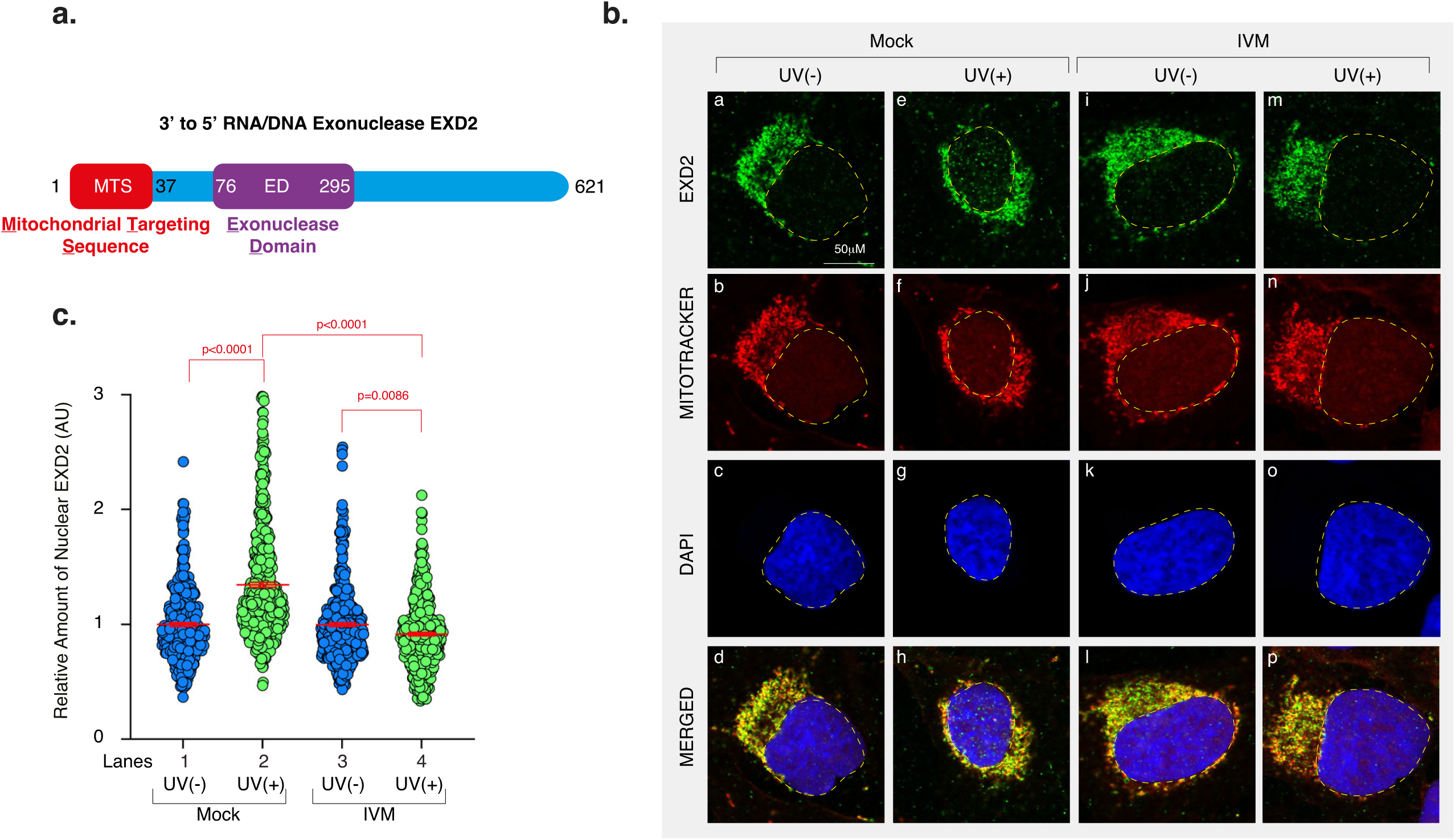
UV irradiation triggers Importin-α/β1-dependent nuclear accumulation of EXD2 (a), Domain architecture of EXD2. EXD2 contains a Mitochondrial Targeting Sequence (MTS) and an Exonuclease Domain (ED). **(b),** Representative confocal images of U-2 OS cells mock or UV-irradiated at 20 J m^-2^. When indicated cells were treated for 1 h with IVM (1μM) before UV irradiation. During the 2 h recovery period, cells were stained with MitoTracker, after which they were fixed at the end of the recovery and labeled with anti-EXD2. MERGED: overlay of EXD2 labeling, MitoTracker and DAPI staining. **(c),** Quantification of EXD2(WT) in the nucleus of U-2 OS cells treated as indicated in **(b)**. Data were normalized to mock-irradiated conditions (set to 1). Red bars indicate mean integrated densities ± s.e.m (*n*>300 cells from three independent experiments). Statistical analyses were performed using one-way ANOVA with Tukey’s post hoc test.

To determine whether EXD2 undergoes stress-induced subcellular relocalization, we monitored endogenous EXD2 distribution by confocal microscopy before and after UV irradiation. In untreated cells, EXD2 displayed a strictly mitochondrial pattern, with nearly complete colocalization between EXD2 staining and MitoTracker (Fig. 2b, panels a-d). Following UV irradiation, a robust nuclear accumulation of endogenous EXD2 was observed (Fig. 2b, panels e-h), which was abolished in IVM-treated cells (Fig. 2b, panels i-p). Quantification confirmed a significant IVM-sensitive increase in nuclear EXD2 levels, with a peak at 2 h post-irradiation (Fig. 2c). These results suggest that EXD2 undergoes UV-induced, Importin-α/β1-dependent import to the nucleus.

### Nuclear-imported EXD2 associates with elongation-arrested RNAPII after UV irradiation

We previously observed an interaction between EXD2 and RNAPII after UV-irradiation (*17*). We next assessed whether this interaction required nuclear import, using proximity ligation assays (PLA). In untreated U-2 OS cells, only low levels of EXD2–RNAPII PLA foci were detected, indicating minimal proximity under basal conditions (Fig. 3a, panels a–d). As previously observed, UV irradiation induced a robust increase in EXD2:RNAPII PLA signals (Fig. 3a, panels e–f and Fig. 3b). Treatment with IVM completely abolished the UV-induced increase in PLA signals, reducing the number of EXD2:RNAPII foci to levels indistinguishable from mock-irradiated cells (Fig. 3a, panels g–h and Fig. 3b). Similarly, the CDK9 inhibitor DRB almost completely prevented the formation of UV-induced EXD2:RNAPII PLA foci (Fig. 3c, d), suggesting that nuclear import and ongoing transcription are necessary for efficient recruitment of EXD2 to RNAPII.

**Fig. 3.**
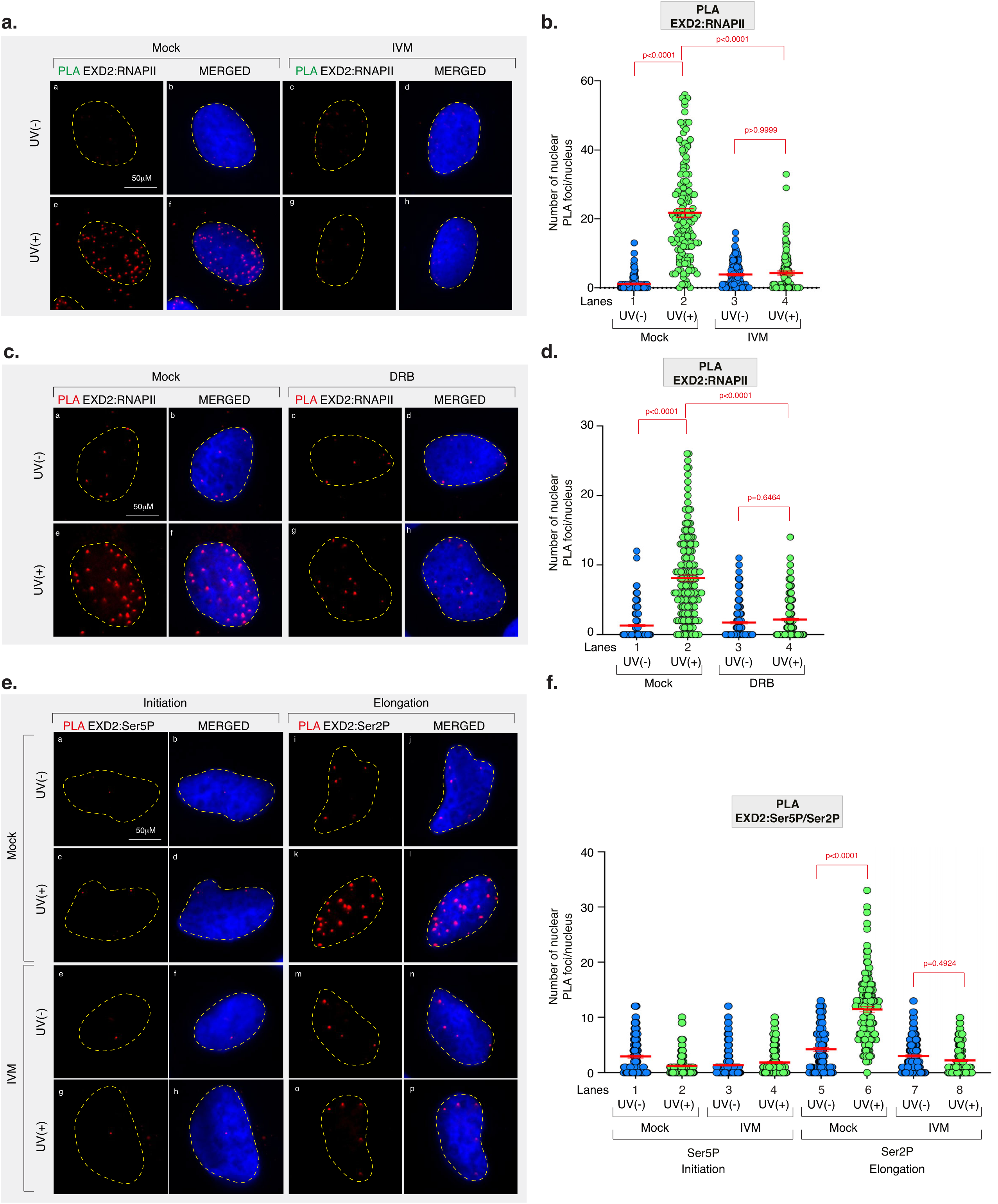
EXD2 interacts with elongation-arrested RNAPII after UV irradiation (a), Representative confocal images of PLA EXD2:RNAPII signal in U-2 OS cells mock-or UV-irradiated at 20 J m^-2^ (recovery time; 2 h). When indicated, cells were treated for 1 h with IVM (1 μM) before UV irradiation. PLA assay was then performed using anti-EXD2 and anti-RNAPII antibodies. MERGED: overlay of PLA and DAPI staining. **(b),** Number of nuclear EXD2:RNAPII PLA foci in U-2 OS cells as described in **(a)**. Red bars indicate mean number of foci ± s.e.m (*n*> 100 cells per condition from three independent experiments). **(c),** Representative confocal images of EXD2:RNAPII PLA signal in U-2 OS cells mock-or UV-irradiated at 20 J m^-2^ (recovery time; 2 h). When indicated, cells were treated for 1 h with DRB (100 μM) before UV irradiation. PLA assay was then performed using anti-EXD2 and anti-RNAPII antibodies. MERGED: overlay of PLA and DAPI staining. **(d),** Number of nuclear EXD2:RNAPII PLA foci in U-2 OS cells as described in **(c)**. Red bars indicate mean number of foci ± s.e.m (*n*> 100 cells per condition from three independent experiments). **(e),** Representative confocal images of EXD2:RNAPII-Ser5P/Ser2P PLA signal in U-2 OS cells mock-or UV-irradiated at 20 J m^-2^ (recovery time; 2 h). When indicated, cells were treated for 1 h with IVM before UV irradiation. PLA assay was then performed using anti-EXD2 and either anti-Ser5P or Ser2P antibodies. MERGED: overlay of PLA and DAPI staining. **(f),** Number of nuclear EXD2:RNAPII-Ser5P/Ser2P PLA foci in U-2 OS cells as described in **(e)**. Red bars indicate mean number of foci ± s.e.m (*n*> 100 cells per condition from three independent experiments). Statistical analyses were performed using one-way ANOVA with Tukey’s post hoc test.

To define which RNAPII species engages EXD2, we combined PLA with antibodies against EXD2 and either Ser5-phosphorylated RNAPII (Ser5P, initiating form) or Ser2-phosphorylated RNAPII (Ser2P, elongating form). Under basal conditions, EXD2:Ser5P and EXD2:Ser2P PLA signals were low (Fig. 3e, panels a-b, i-j and Fig. 3f). Following UV irradiation, EXD2:Ser2P PLA foci increased markedly in mock-treated cells, whereas EXD2:Ser5P PLA signals were not affected (Fig. 3e, panels c-d, k-l and Fig. 3f). Importantly, IVM treatment abolished the UV-induced increase in EXD2:Ser2P PLA foci (Fig. 3e, panels e-h, m-p and Fig. 3f).

Collectively, these data indicate that EXD2 is in close proximity to elongation-arrested RNAPII following UV damage, and that this proximity requires its stress-induced nuclear import.

### A functional NLS is required for stress-induced nuclear import of EXD2

Prompted by the above findings, we investigated whether EXD2 contains a functional nuclear localization sequence (NLS). Using cNLS Mapper (*38*), we identified a putative NLS located from amino-acid 364 to 374 (Fig. 4a). To determine whether the predicted NLS is required for the UV-induced nuclear relocalization of EXD2, we generated U-2 OS cell lines in which GFP was knocked-in at the endogenous EXD2 locus (EXD2(WT)::GFP) and produced two clones in which the NLS was mutated (EXD2(mutNLS)::GFP, clones 1 and 2) (Fig. 4b).

**Fig. 4.**
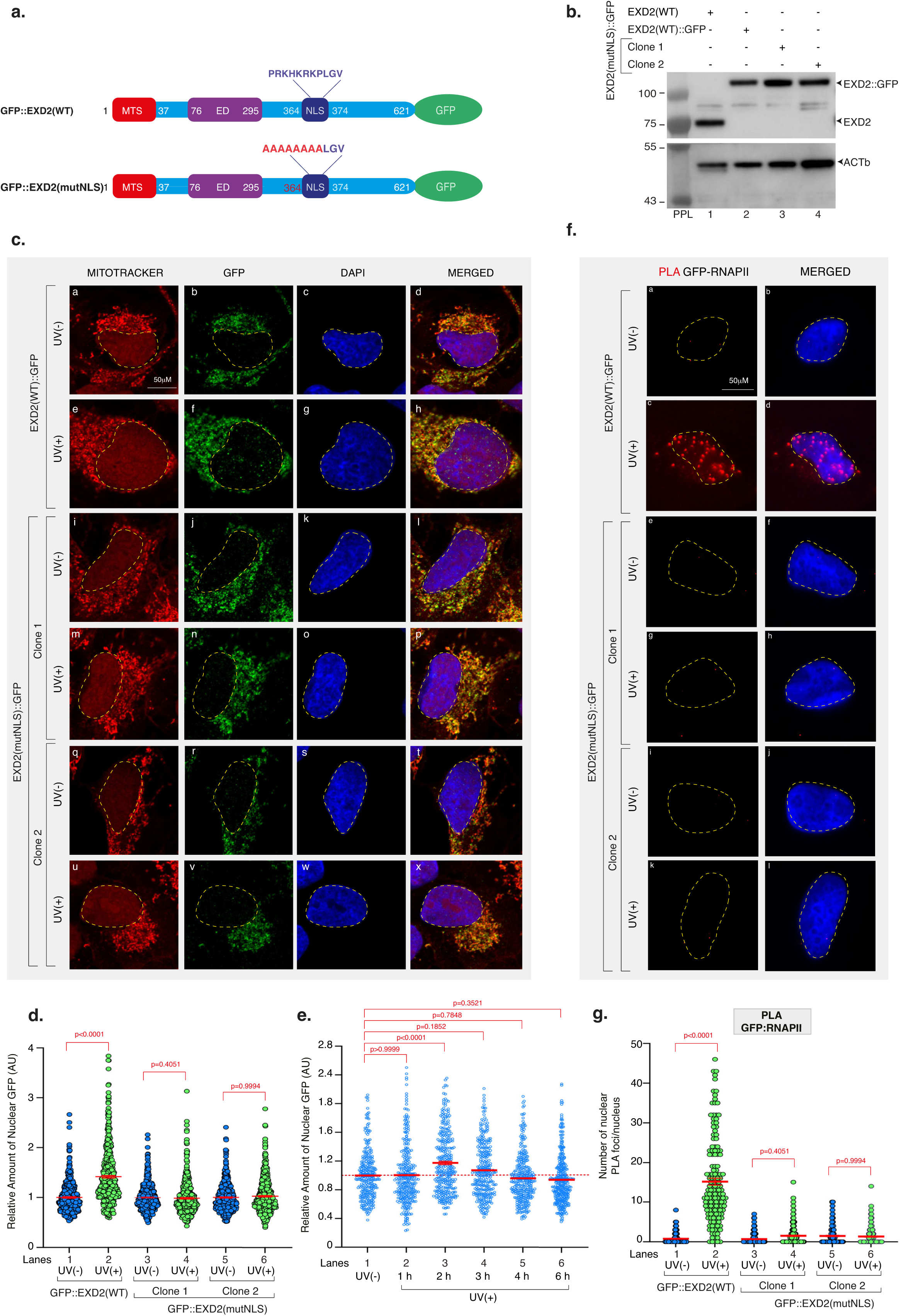
Mutation of the EXD2 NLS abolishes its UV-induced nuclear import (a), Domain architecture of the EXD2(WT)::GFP and EXD2(mutNLS)::GFP genes in CRISPR/Cas9 engineered U-2 OS cells. Sequence of the NLS (dark blue) is indicated. In red the residues mutated in U-2 OS EXD2(mutNLS)::GFP cells are indicated. **(b),** Protein lysates of U-2 OS EXD2(WT), EXD2(WT)::GFP and EXD2(mutNLS)::GFP (clones 1 and 2) cells were immuno-blotted for EXD2 or ACTb as indicated. Molecular weights (kDa) and pre-stained protein ladders (PPL) are shown. **(c),** Representative confocal images of U-2 OS EXD2(WT)::GFP and U-2 OS EXD2(mutNLS)::GFP (clones 1 and 2) cells mock-or UV-irradiated at 20 J m^-2^ (recovery time; 2 h). Cells were stained with MitoTracker, after which they were fixed at the end of the recovery and labeled with anti-GFP. MERGED: overlay of GFP labeling, MitoTracker and DAPI staining. **(d),** Quantification of EXD2(WT)::GFP and EXD2(mutNLS)::GFP (in clones 1 and 2) in the nucleus of U-2 OS cells treated as indicated in **(c)**. Data were normalized to the mock treatment controls (set to 1). Red bars indicate mean integrated densities ± s.e.m (*n* > 400 cells per condition from three independent experiments). **(e),** Quantification of EXD2(WT)::GFP in the nucleus of U-2 OS cells mock-or UV-irradiated (20 J m^-2^) and fixed at the indicated recovery times. Data were normalized to the mock treatment controls (set to 1). Red bars indicate mean integrated densities ± s.e.m (n> 100 cells from three independent experiments). **(f),** Representative confocal images of PLA GFP-RNAPII signal in U-2 OS EXD2(WT)::GFP and EXD2(mutNLS)::GFP (clones 1 and 2) cells mock-or UV-irradiated at 20 J m^-2^ (recovery time; 2 h). PLA assay was performed using anti-GFP and anti-RNAPII antibodies. MERGED: overlay of PLA and DAPI staining. **(g),** Number of nuclear GFP/RNAPII PLA foci in cells treated like in **(f)**. Red bars indicate mean number of foci ± s.e.m (*n*> 100 cells per condition from three independent experiments). Statistical analyses were performed using one-way ANOVA with Tukey’s post hoc test.

Consistent with our observations for endogenous EXD2 in wild-type cells, confocal microscopy of EXD2(WT)::GFP cells revealed a predominantly mitochondrial GFP signal under non-stressed conditions (Fig. 4c, panels a–d). UV irradiation induced a clear increase in nuclear EXD2(WT)::GFP (Fig. 4c, panels e–h), corresponding to a ∼15–20% elevation in nuclear EXD2 levels (Fig. 4d). In striking contrast, in both EXD2(mutNLS)::GFP clones, EXD2 failed to be imported to the nucleus after UV irradiation (Fig. 4c, panels i–x and Fig. 4d), demonstrating that the NLS is essential for stress-induced nuclear import. A time-course analysis confirmed the transient nature of EXD2(WT)::GFP nuclear accumulation, which peaked at 2 h and returned toward baseline by 4–6 h after UV exposure (Fig. 4e).

We next asked whether the absence of NLS impairs the interaction of EXD2 with RNAPII. PLA analysis using anti-GFP and anti-RNAPII antibodies showed a strong UV-dependent accumulation of PLA foci in EXD2(WT)::GFP cells (Fig. 4f, panels a–d and Fig. 4g). This UV-induced proximity was completely abolished in both EXD2(mutNLS)::GFP clones (Fig. 4f, panels e–l and Fig. 4g). We conclude from these experiments that NLS-mediated EXD2 import is required for its proximity to elongation-arrested RNAPII following UV irradiation.

### EXD2 NLS is essential for transcription recovery and UV resistance

We then assessed how loss of EXD2 nuclear import affects transcription recovery after genotoxic stress. In contrast to EXD2(WT)::GFP cells, EXD2(mutNLS)::GFP cells failed to restore transcription, exhibiting a complete block in RRS similar to the phenotype observed upon global inhibition of nuclear import (Fig. 5a-b). Furthermore, both EXD2(mutNLS)::GFP clones displayed pronounced UV hypersensitivity compared to EXD2(WT)::GFP (Fig. 5c), consistent with their failure to restore transcription. Given the involvement of EXD2 in HRR ^31, 32^, we also tested whether this repair pathway was affected in EXD2(mutNLS)::GFP cells. Using a plasmid-based HRR assay, we found that HRR activity was significantly reduced in EXD2(mutNLS)::GFP cells, and that IVM treatment reproduced this defect in EXD2(WT)::GFP cells, strongly suggesting that HRR also requires the Importin-α/β1-dependent nuclear import of EXD2 (Fig. 5d).

**Fig. 5.**
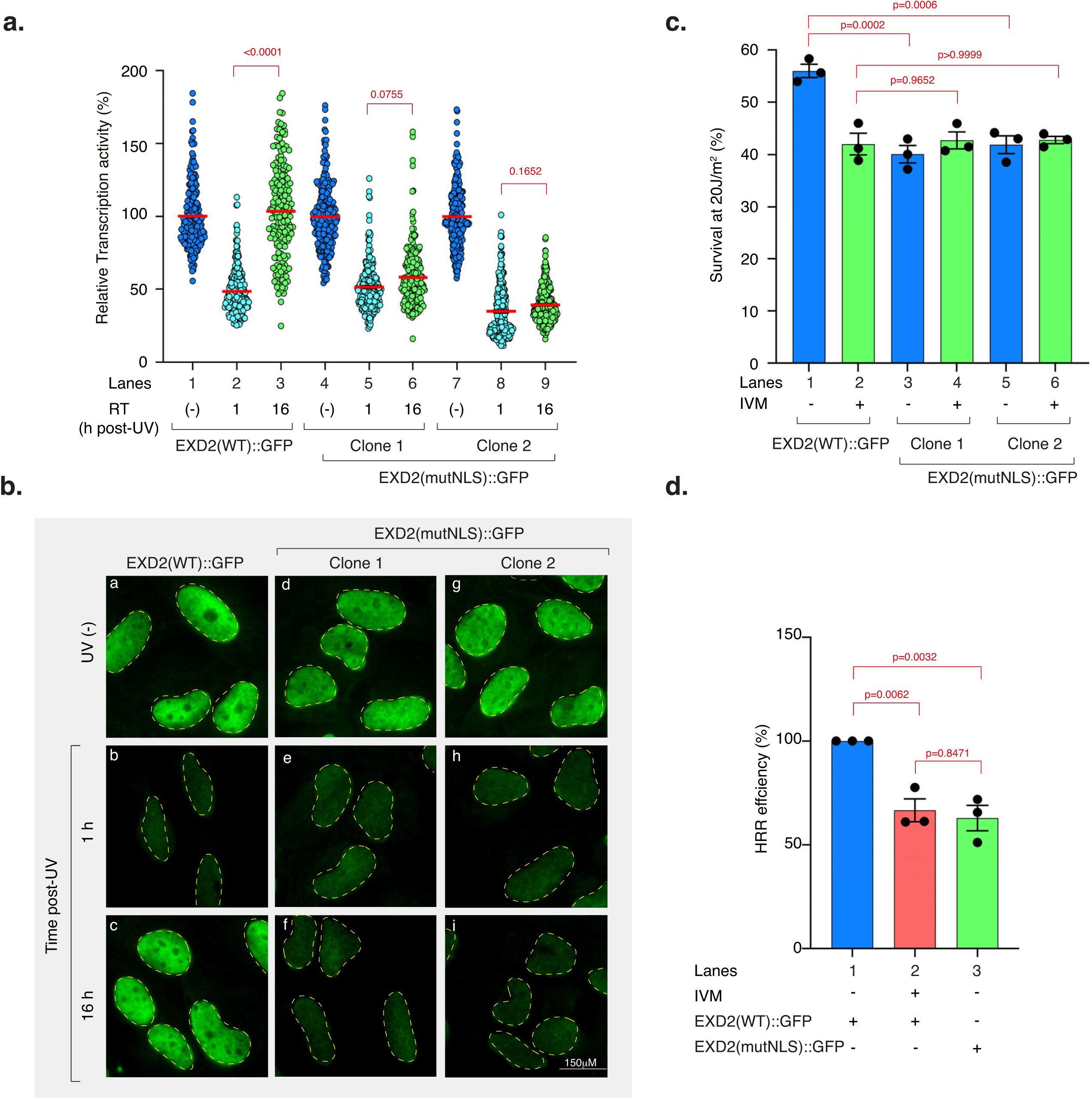
Mutation of the EXD2 NLS blocks transcription recovery and reduces UV resistance (a), Transcription rate in U-2 OS EXD2(WT)::GFP and U-2 OS EXD2(mutNLS)::GFP (clones 1 and 2) cells mock-or UV-irradiated at 20 J m^-2^. mRNA was labeled with EU at the indicated time points post-UV. To ensure specificity for RNAPII, cells were pre-treated with a low dose of actinomycin D to inhibit RNAPI-driven transcription. EU signal was quantified by Fiji (ImageJ) and relative integrated densities, normalized to mock-irradiated level set to 100%, are reported on the graph. Red bars indicate mean integrated densities ± s.e.m (*n*> 200 cells per condition from three independent experiments). RT; recovery time. (-); cells were mock-irradiated. **(b),** Representative microscopy images of U-2 OS EXD2(WT)::GFP and U-2 OS EXD2(mutNLS)::GFP (clones 1 and 2) cells treated like in **(a)**. **(c),** Cell survival of U-2 OS EXD2(WT)::GFP and U-2 OS EXD2(mutNLS)::GFP (clones 1 and 2) cells treated with UV irradiation (20 J m^-2^). Cells were allowed to recover for 48 h and survival was determined. Data were normalized to non-irradiated controls (set to 100%). Data are mean ± s.e.m. from three independent experiments. **(d),** Quantification of HRR pathway in U-2 OS EXD2(WT)::GFP and U-2 OS EXD2(mutNLS)::GFP (clone 1) cells. When indicated, cells were treated for 1 h with IVM (1 μM) before HRR assay plasmids transfection. Data were normalized to the mock-treated U-2 OS EXD2(WT)::GFP cells (set to 100%). Data are mean ± s.e.m. from three independent experiments. Statistical analyses were performed using one-way ANOVA with Tukey’s post hoc test.

Collectively, these data demonstrate that the EXD2 NLS is indispensable for its function in cell survival following genotoxic stress. Mutation of the NLS phenocopies the pharmacological inhibition of Importin-α/β1-dependent nuclear import, confirming that stress-induced nuclear import of EXD2 is a critical mechanistic step in transcription recovery, and more broadly in the nuclear functions of EXD2.

### Constitutive nuclear EXD2 engages with RNAPII after UV, independently of nuclear import

To determine whether EXD2 nuclear import is the limiting event enabling its association with RNAPII after UV irradiation, we assessed the behavior of EXD2 variants that are constitutively nuclear. For this purpose, we used HeLa cells in which endogenous EXD2 was either intact, deleted, or complemented with FLAG-HA–tagged versions of EXD2 lacking the MTS (*29*), (*30*) (Fig. 6a-b). As expected, wild-type EXD2 in HeLa EXD2(+/+) cells localized almost exclusively to mitochondria under basal conditions, while EXD2 was undetectable in EXD2(−/−) cells (Fig. 6c, panels a-h). Expression of an EXD2 mutant lacking the MTS (EXD2(ΔMTS)) redistributed a fraction of the protein to the nucleus, confirming that removal of the MTS is sufficient to bypass mitochondrial sequestration (Fig. 6c, panels i-l). A nuclease-dead version of EXD2(ΔMTS) carrying D108A/E110A substitutions (EXD2(ΔMTS/ND)) (*32*) displayed an identical nuclear pattern (Fig. 6a-b and Fig. 6c, panels m-p).

**Fig. 6.**
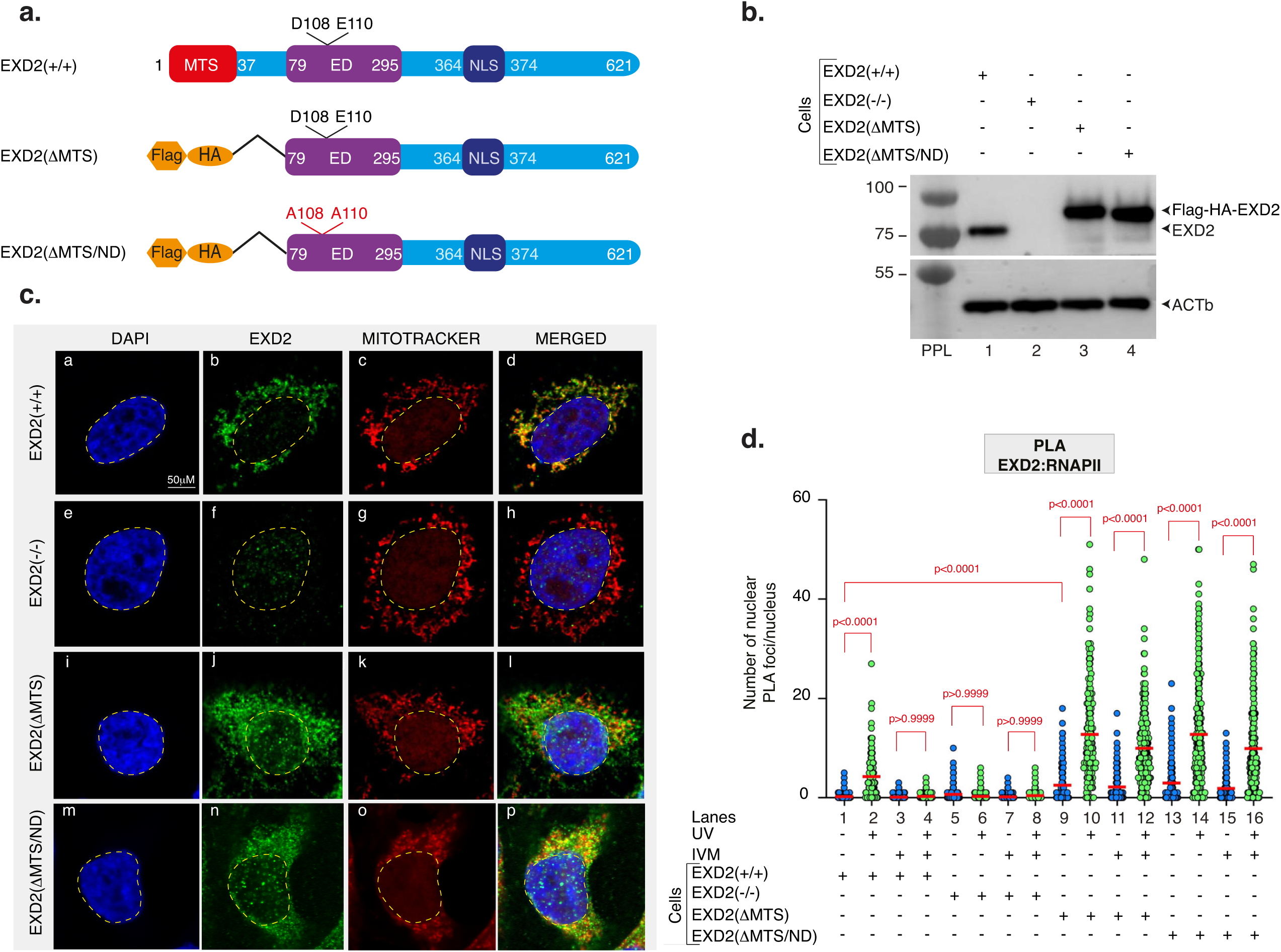
Forced nuclear EXD2 interacts with RNAPII after UV independently of nuclear import (a), Domain architecture of either the endogenous EXD2(+/+) in HeLa cells or the FLAG-HA-EXD2(ΔMTS) and FLAG-HA-EXD2(ΔMTS/ND) constructs stably expressed in HeLa EXD2(-/-) cells (EXD2(ΔMTS) and EXD2(ΔMTS/ND) cells respectively). **(b),** Protein lysates of HeLa EXD2(+/+), EXD2(-/-), EXD2(ΔMTS) and EXD2(ΔMTS/ND) cells were immuno-blotted for EXD2 or ACTb as indicated. Molecular weights (kDa) and prestained protein ladders are shown (PPL) are shown. **(c),** Representative confocal images of HeLa EXD2(+/+), EXD2(-/-), EXD2(ΔMTS) and EXD2(ΔMTS/ND). Cells were stained with MitoTracker and labeled with anti-EXD2. MERGED: overlay of EXD2 labeling, MitoTracker and DAPI staining. **(d),** Number of nuclear EXD2:RNAPII PLA foci in mock-or UV-irradiated (20 J m^-2^, recovery time; 2 h) HeLa cells as indicated. When indicated, cells were treated for 1 h with IVM (1 μM) before UV irradiation. PLA assay was then performed using anti-EXD2 and anti-RNAPII antibodies. Red bars indicate mean number of foci ± s.e.m (*n*> 100 cells per condition from three independent experiments). Statistical analyses were performed using one-way ANOVA with Tukey’s post hoc test.

We next examined whether forced nuclear EXD2 is able to associate with RNAPII. As observed above in U-2 OS cells, a robust UV-induced and IVM-sensitive increase in EXD2:RNAPII PLA signals was detected in wild-type HeLa cells (Fig. 6d, lanes 1-4).

As expected, no PLA signal was observed in EXD2(−/−) cells (lanes 5-8). Strikingly, expression of EXD2(ΔMTS) in HeLa EXD2(-/-) cells restored the UV-induced EXD2:RNAPII PLA signal, but this signal could now also be detected in the presence of IVM (lanes 9–12). Notably, EXD2(ΔMTS) also showed a low but significant proximity with RNAPII in unstressed cells (compare lane 1 with lane 9). The nuclease-dead EXD2(ΔMTS/ND) mutant displayed similar behavior, retaining RNAPII proximity both before and after UV regardless of IVM treatment, with a significant increase after UV exposure (lanes 13–16).

These results show that constitutive nuclear targeting of EXD2 is sufficient to restore its UV-induced proximity with RNAPII even when nuclear import is pharmacologically blocked. This suggests that the critical function of Importin-α/β1–mediated import is to supply a nuclear pool of EXD2, and that once in the nucleus, this pool is intrinsically competent to engage elongating RNAPII, predominantly after UV irradiation.

### Constitutive nuclear EXD2 restores RRS and UV resistance but induces genome instability

We next examined transcription recovery following UV exposure in HeLa cells used above. EXD2 depletion or expression of the nuclease-dead mutant impaired RRS under both basal and IVM-treated conditions, compared to EXD2(+/+) (Fig. 7a, lanes 4-6, 10-12, 16-18 and 22-24 and Fig. 7b, panels g-l and s-x). Remarkably, expression of constitutively nuclear EXD2(ΔMTS) fully restored RRS, even in IVM-treated cells (Fig. 7a, lanes 7-9 and lanes 19-21 and Fig. 7b, panels m-r).

**Fig. 7.**
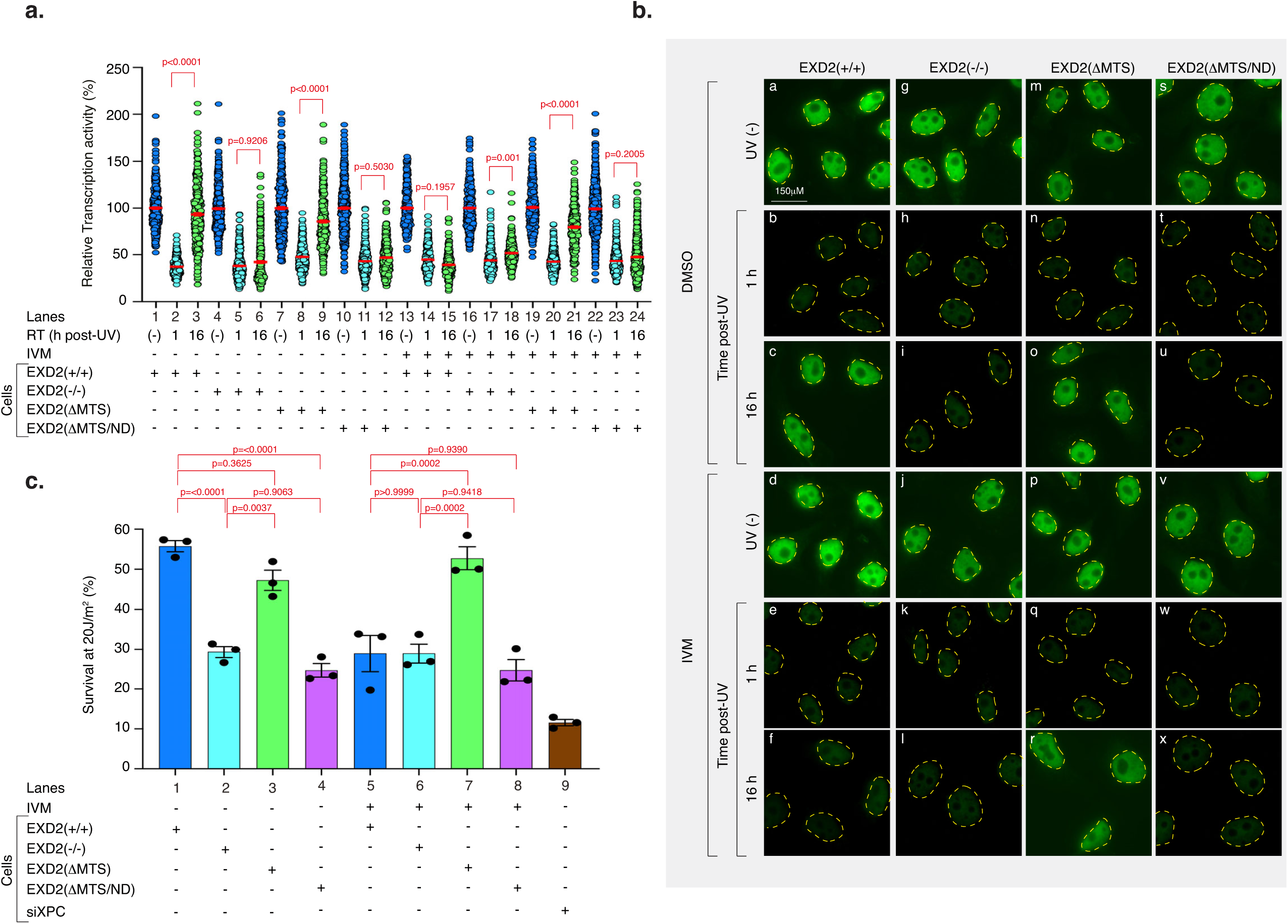
Nuclear-targeted EXD2 rescues RRS and UV resistance **(a),** Transcription rate in HeLa EXD2(+/+), EXD2(-/-), EXD2(ΔMTS), and EXD2(ΔMTS/ND) cells mock-or UV-irradiated at 20 J m^-2^. mRNA was labeled with EU at the indicated time points post-UV. To ensure specificity for RNAPII, cells were pre-treated with a low dose of actinomycin D to inhibit RNAPI-driven transcription. When indicated, cells were treated for 1 h with IVM (1 μM) before UV irradiation. EU signal was quantified by Fiji (ImageJ) and relative integrated densities, normalized to mock-treated level set to 100%, are reported on the graph. Red bars indicate mean integrated densities ± s.e.m (*n*> 250 cells per condition from three independent experiments). RT; recovery time. (-); cells were mock-irradiated. **(b),** Representative microscopy images of HeLa cells treated like in **(a)**. **(c),** Cell survival of HeLa EXD2(+/+), EXD2(-/-), EXD2(ΔMTS) and EXD2(ΔMTS/ND) cells treated with UV irradiation (20 J m^-2^). Cells were allowed to recover for 48 h and survival was determined. When indicated, cells were treated for 1 h with IVM (1 μM) or for 48 h with siXPC before UV irradiation Data were normalized to the mock-irradiated controls (set to 100%). Data are mean ± s.e.m from three independent experiments. Statistical analyses were performed using one-way ANOVA with Tukey’s post hoc test.

We then evaluated UV resistance across these cell lines. Without IVM, HeLa EXD2(+/+) and HeLa EXD2(-/-) + EXD2(ΔMTS) cells exhibited comparable UV resistance (Fig. 7c, compare lanes 1 and 3, and Sup. Fig 2, left panel), while HeLa EXD2(-/-) and EXD2(ΔMTS/ND) cells were hypersensitive (Fig. 7c, compare lanes 1, 3 with lanes 2, 4 and Sup. Fig. 2, Left panel). Under IVM treatment, HeLa EXD2(+/+) cells became as sensitive as HeLa EXD2(-/-) and HeLa EXD2(-/-) + EXD2(ΔMTS/ND) cells (Fig. 7c, compare lane 5 with lanes 6, 8, and Sup. Fig. 2, right panel). In marked contrast, HeLa EXD2(-/-) + EXD2(ΔMTS) cells retained robust resistance regardless of IVM treatment (Fig. 7c, compare lane 3 with lane 7 and Sup. Fig 2, right panel).

We next investigated whether constitutive nuclear EXD2 could lead to abnormal mitotic phenotypes. Immunofluorescence analysis revealed that HeLa EXD2(-/-) + EXD2(ΔMTS) cells displayed numerous mitotic defects, including misaligned chromosomes at metaphase, less frequently observed in HeLa EXD2(+/+) or EXD2(-/-) cells (Sup. Fig. 3a, b). The nuclease-dead EXD2 mutant expressed in HeLa EXD2(-/-) + EXD2(ΔMTS/ND) cells led to a normal mitotic profile, implicating EXD2’s nuclease activity in the observed mitotic defect. We then examined the consequences of these alignment defects on genome stability by quantifying micronuclei, which arise from lagging chromosomes or chromosome fragments that fail to be incorporated into daughter nuclei (*39*). EXD2(+/+) cells showed detectable basal micronuclei levels (Sup. Fig. 3c, d). EXD2(-/-) cells displayed a modest increase, consistent with a general role of EXD2 in safeguarding genome integrity. Strikingly, EXD2(ΔMTS) cells exhibited a robust accumulation of micronuclei, by far the highest among all genotypes examined, indicating frequent chromosome segregation errors. As for metaphase misalignments, the EXD2(ΔMTS/ND) mutant significantly reduced micronuclei formation, confirming that excessive, unregulated nuclear EXD2 nuclease activity drives the observed mitotic defects and genomic instability.

Together, these data show that constitutively nuclear EXD2 is sufficient to restore transcription restart and UV resistance in EXD2-deficient cells. However, the persistence of its exonuclease activity in the nucleus disrupts mitotic chromosome alignment and promotes micronuclei formation.

### EXD2 nuclear import licenses resolution of UV-induced RNA–DNA hybrids

Our data support a model in which stress-induced nuclear import of EXD2 limits the accumulation of elongation-arrested RNAPII complexes associated with R-loop-like RNA–DNA hybrid structures (hereafter referred to as R-loop–like intermediates). To visualize R-loop–like intermediates, we used a catalytically inactive RNaseH1 mutant (dRNaseH1-GFP), which binds RNA–DNA hybrids without degrading them and has been widely used as a specific sensor of R-loops (*40*), (*41*). To determine whether RNAPII associates with R-loop–like intermediates in UV-irradiated cells, we assessed the proximity of dRNaseH1-GFP to RNAPII by PLA. A robust PLA signal between RNAPII and dRNaseH1-GFP was detected 2 h after UV irradiation and was abolished by transcription inhibition with DRB (Fig. 8a). Similarly, EXD2 displayed UV-induced proximity to dRNaseH1-GFP 2 h after irradiation, which was abolished by DRB (Fig 8b). We next investigated whether the association between RNAPII complexes and R-loop–like intermediates was modulated by EXD2. We observed that the PLA signal between RNAPII and dRNaseH1-GFP was robust 2 h after UV irradiation but disappeared 16 h later in HeLa EXD2(+/+) cells. In contrast, in HeLa EXD2(−/−) cells, the PLA signal strongly persisted at 16 h post-irradiation. (Fig. 8c and Sup. Fig. 4a). These results suggest that EXD2 is required to license the resolution of transcription-associated R-loop–like intermediates formed at elongation-arrested RNAPII complexes following UV irradiation.

**Fig. 8.**
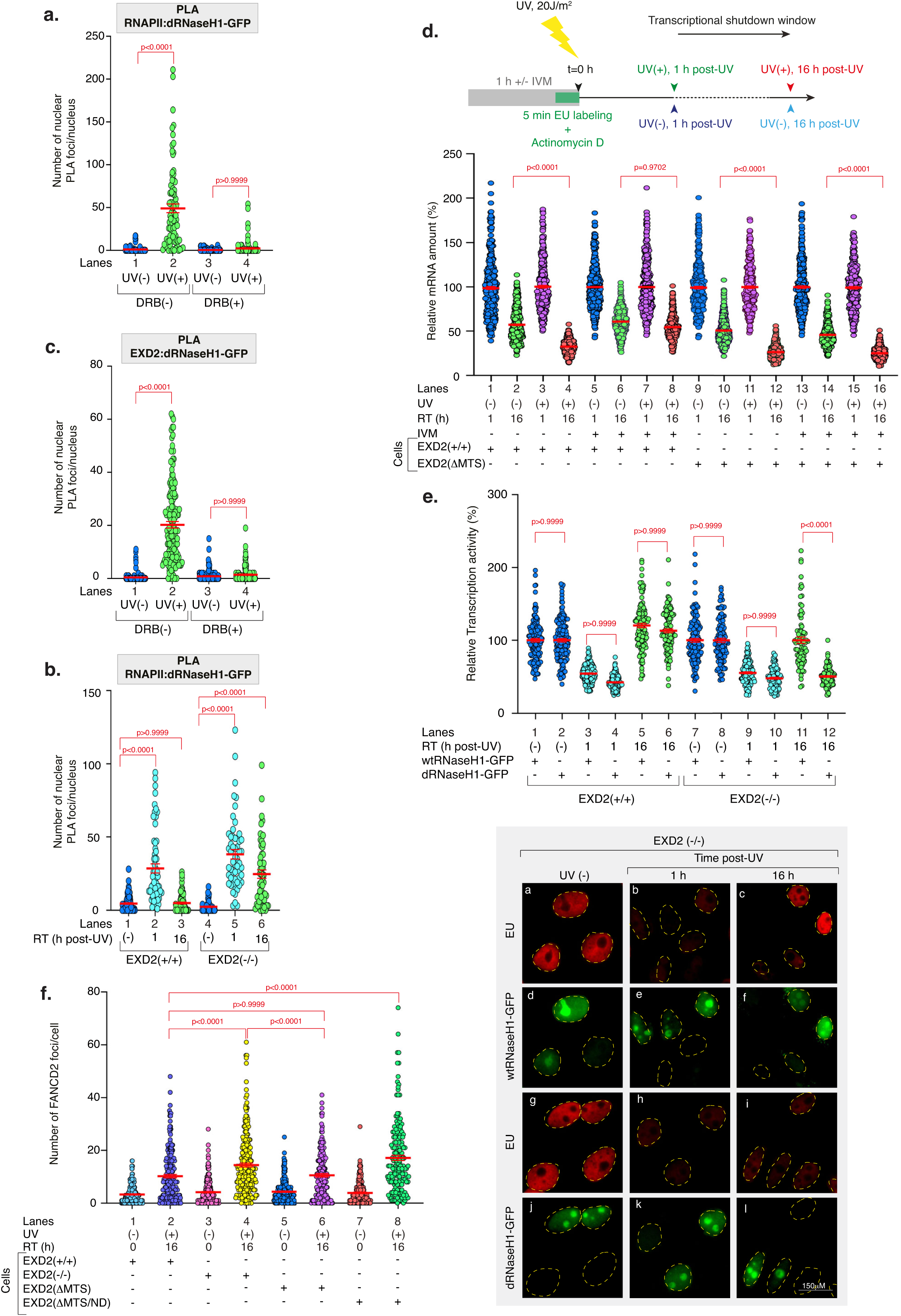
EXD2 resolves UV-induced and RNAPII-associated R-loop-like intermediates (a), Number of nuclear RNAPII:dRNaseH1-GFP PLA foci in mock-or UV-irradiated (20 J m^-2^, recovery time; 2 h) U-2 OS cells. When indicated, cells were treated for 1 h with DRB (100 μM) before UV irradiation to inhibit transcription. PLA assay was then performed using anti-GFP and anti-RNAPII antibodies. Red bars indicate mean number of foci ± s.e.m (*n*> 100 cells per condition from three independent experiments). **(b),** Number of nuclear RNAPII:dRNaseH1-GFP PLA foci in mock-or UV-irradiated (20 J m^-2^) HeLa EXD2(+/+) and EXD2(-/-) cells at the indicated time points post-UV. PLA assay was performed using anti-GFP and anti-RNAPII antibodies. Red bars indicate mean number of foci ± s.e.m (*n*> 100 cells per condition from three independent experiments). **(c),** Number of nuclear flag-EXD2:dRNaseH1-GFP PLA foci in mock-or UV-irradiated (20 J m^-2^, recovery time; 2 h) in HeLa EXD2 (ΔMTS) cells. When indicated, cells were treated for 1 h with DRB before UV irradiation to inhibit transcription. PLA assay was performed using anti-GFP and anti-flag antibodies. Red bars indicate mean number of foci ± s.e.m (*n*> 100 cells per condition from three independent experiments). **(d), Upper panel;** Scheme of the EU pulse-chase assay. Cells were incubated for 1 h with DMSO or IVM (1 μM), for 30 min with Actinomycin D (0.05μg mL^-1^) to specifically inhibit RNAPI transcription and mRNAs were pulse-labeled with EU for 10 min prior to UV irradiation (20 J m^-2^). Cells were allowed to recover for 1 h or 16 h post-UV before fixation and EU intensity was measured using Fiji (ImageJ). Actinomycin D was maintained during the experiment. **Lower panel;** HeLa EXD2(+/+) and EXD2(ΔMTS/ND) cells were treated as above and EU signals were quantified using Fiji (ImageJ) and normalized to the value obtained at 1 h set to 100%. Values are reported on the graph. Red bars indicate mean integrated densities ± s.e.m (*n*> 300 cells per condition from three independent experiments). RT; recovery time. **(e),** Transcription rate in HeLa EXD2(+/+) and EXD2(-/-) cells expressing either wtRNaseH1-GFP or an inactive dRNaseH1-GFP (D210N) mock-or UV-irradiated at 20 J m^-2^. mRNA was labeled with EU at the indicated time points post-UV. To ensure specificity for RNAPII, cells were pre-treated with a low dose of actinomycin D to inhibit RNAPI-driven transcription. **Upper panel**; EU signal was quantified by Fiji (ImageJ) and relative integrated densities, normalized to mock-treated level set to 100%, are reported on the graph. Red bars indicate mean integrated densities ± s.e.m (*n*> 300 cells per condition from three independent experiments). RT; recovery time. (-); cells were mock-irradiated. **Lower panel**; Representative microscopy images of HeLa EXD2(-/-) cells expressing either wtRNaseH1-GFP or an inactive dRNaseH1-GFP (D210N) and treated as described above. **(f),** Quantification of FANCD2 foci per nucleus in HeLa EXD2(+/+), EXD2(-/-), EXD2(ΔMTS), and EXD2(ΔMTS/ND) cells mock or UV-irradiated at 20 J m^-2^ (recovery time; 16h). After recovery time, cells were fixed, labeled with anti-FANCD2 and foci were quantified by Fiji (ImageJ) and reported on the graph. Red bars indicate mean number of foci ± s.e.m (*n*> 200 cells per condition from three independent experiments). RT; Recovery time. Statistical analyses were performed using one-way ANOVA with Tukey’s post hoc test.

EXD2 possesses 3′–5′ exonuclease activity with a preference for degrading the RNA strand of RNA–DNA hybrids *in vitro* (*42*). We therefore asked whether EXD2 promotes degradation of nascent RNA existing in UV-irradiated cells and which could be part of the R-loop–like intermediates when RNAPII is blocked during elongation. To test this hypothesis, nascent transcripts were pulse-labeled by a short EU incubation immediately before UV irradiation, while RNAPI transcription was inhibited. EU-labeled mRNA was monitored after 1 h and 16 h to assess degradation of transcripts synthesized just before transcriptional shutdown (Fig. 8d). In mock-irradiated HeLa EXD2(+/+) and EXD2(-/-)+EXD2(ΔMTS) cells, EU-labeled RNA declined by ∼40–50% between 1 h and 16 h, irrespective of IVM treatment, consistent with basal mRNA turnover (*43*) (Fig. 8d, lanes 1–2, 5–6, 9–10, 13–14). UV irradiation increased RNA loss to ∼70% in the absence of IVM (Fig. 8d, lanes 3–4, 11–12), consistent with enhanced degradation of RNA associated with stalled transcription complexes. Strikingly, blocking EXD2 nuclear import with IVM reduced this UV-induced RNA degradation in EXD2(+/+) cells to near-baseline levels (Fig. 8d, lanes 7–8), whereas cells expressing constitutively nuclear EXD2(ΔMTS) retained the enhanced RNA degradation despite IVM treatment (Fig. 8d, lanes 15–16). These findings suggest a direct role for EXD2 nuclease activity in degrading RNA within R-loop–like intermediates formed at elongation-arrested RNAPII and demonstrate the central role of nuclear import in RNA–DNA metabolism post-UV irradiation.

Because resolution of R-loop-like RNA–DNA hybrids primarily depends on RNaseH activity (*44*), we next tested whether RNaseH1 overexpression could compensate for EXD2 loss. Overexpression of wild-type RNaseH1 (wtRNaseH1-GFP) in HeLa (EXD2-/-) cells restored transcription recovery 16 h after UV irradiation, whereas dRNaseH1-GFP failed to do so (Fig. 8e, upper panel). Notably, fluorescence microscopy indicated that transcription is restored specifically in cells expressing wtRNaseH1-GFP (Fig. 8e, lower panel). Overexpression of wtRNaseH1-GFP or dRNaseH1-GFP did not modify transcription recovery in HeLa (EXD2+/+) cells (Fig. 8e, upper panel and Sup. Fig. 4b). These data support the idea that EXD2 promotes transcription recovery by facilitating resolution of R-loop–like intermediates formed during transcription shutdown.

We then asked whether defective RNA processing in EXD2-deficient cells leads to TRCs. Quantification of FANCD2 foci, a marker of replication stress frequently associated with TRCs (*45*), revealed a marked increase 16 h after UV irradiation in EXD2-knockout cells compared with wild type (Fig. 8f and Sup. Fig. 4c). Expression of constitutively nuclear EXD2(ΔMTS) restored FANCD2 foci to near-wild-type levels, whereas the nuclease-dead EXD2 mutant failed to do so, demonstrating that EXD2 nuclease activity is required to resolve toxic TRCs in stressed cells.

Finally, to extend our analysis to additional genotoxic stresses, we tested whether pharmacological inhibition of EXD2 nuclear import could enhance the efficacy of transcription-blocking chemotherapeutic agents. Synergy analysis using the Highest Single Agent (HSA) model revealed strong synergistic cytotoxicity between IVM and cisplatin in HeLa EXD2(+/+) cells (HSA score 13.02; Sup. Fig. 5a). In contrast, synergy was markedly reduced in cells expressing constitutively nuclear EXD2(ΔMTS) (HSA score 4.22; Sup. Figs. 5b, c), indicating that inhibition of EXD2 nuclear import underlies the combinatorial effect.

Collectively, these results indicate that stress-induced nuclear import of EXD2 is a key mechanism that promotes degradation of RNA within R-loop-like intermediate structures formed at stalled RNAPII complexes, thereby limiting TRCs and associated cytotoxicity. They also suggest that this nuclear deployment is a general response to genotoxic stress, which could be targeted to enhance the efficacy of transcription-blocking chemotherapeutic agents.

## Discussion

Cells must efficiently coordinate DNA repair with the restoration of transcription following genotoxic stress. While transcriptional shutdown after DNA damage has been extensively characterized, the mechanisms that enable transcription restart once lesions are removed remain incompletely understood. Our findings identify regulated nuclear import of the RNA/DNA exonuclease EXD2 as a key step in this process and suggest that spatial control of nuclease availability represents an additional level of regulation within the DDR. We propose that stress-induced nuclear import of EXD2 functions as a licensing step that transiently enables RNA–DNA hybrid processing at stalled transcription complexes while preventing deleterious nuclease activity under basal conditions.

A central feature of this mechanism is the dynamic subcellular localization of EXD2. Under basal conditions, EXD2 is predominantly mitochondrial, consistent with its established role in mitochondrial genome expression (*29*), (*30*). Following UV-induced genotoxic stress, however, a fraction of EXD2 transiently accumulates in the nucleus in an Importin-α/β1–dependent manner. Preventing this relocalization, either pharmacologically or through mutation of its nuclear localization signal, blocks transcription recovery despite apparently normal removal of UV-induced DNA lesions. Conversely, constitutive nuclear targeting of nuclease-active EXD2 restores transcription restart even when nuclear import is inhibited. Together, these observations indicate that regulated nuclear access of EXD2 represents a limiting step in transcription recovery following DNA damage.

Our results further suggest that EXD2 promotes transcription restart by processing R-loop–like intermediates, consistent with a direct role in RNA–DNA hybrid metabolism, although indirect effects on RNA stability cannot be formally excluded. Increasing evidence indicates that transcriptional arrest can favor the formation of R-loop–like structures, particularly when RNAPII backtracking displaces the nascent RNA from the catalytic center (*46*), (*47*). In such contexts, the extruded RNA strand may rehybridize with the DNA template, generating RNA–DNA hybrid intermediates that can interfere with transcription restart if not properly resolved. Consistent with this model, we observe UV-induced proximity between RNAPII and a catalytically inactive RNaseH1 mutant used as a sensor of RNA–DNA hybrids, as well as between EXD2 and this sensor. Importantly, these interactions are transcription-dependent and require EXD2 nuclear import.

Rather than relying on S9.6 immunodetection, which can recognize additional RNA-containing structures (*48*), (*49*), we used a catalytically inactive RNaseH1 mutant as a sensor of R-loop–like intermediates (*40*), (*41*). The combination of RNaseH1-based detection, transcription dependence, and functional rescue by wild-type RNaseH1 provides convergent evidence supporting the involvement of R-loop–like intermediates in this context. In agreement with this interpretation, EXD2 deficiency leads to persistent RNAPII complexes associated with unresolved R-loop–like intermediates, whereas overexpression of wild-type RNaseH1 restores transcription recovery. Together, these findings support a model in which EXD2 promotes transcription restart by facilitating the resolution of transcription-associated R-loop–like intermediates. The recent demonstration that EXD2 preferentially degrades the RNA strand of R-loops to promote their resolution *in vitro* further supports our model (*42*). We refer to these species as “R-loop–like intermediates” because their precise molecular configuration remains unknown. Consequently, unbiased genome-wide detection remains challenging. Their biochemical and structural characterization will therefore be an important direction for future investigation.

Our data also link defective EXD2 nuclear import with increased replication stress following UV irradiation. We observed that impaired EXD2 nuclear import is associated with elevated FANCD2 foci formation, indicative of increased replication stress and potentially enhanced transcription–replication conflicts (*32*). Although we did not directly visualize transcription–replication collisions, the increase in FANCD2 foci is consistent with replication stress associated with persistent RNA–DNA hybrids, which have been widely linked to genome instability in both eukaryotes and prokaryotes (*20*), (*50*). In this context, our results support a model in which EXD2-mediated processing of transcription-associated R-loop–like intermediates contributes to genome stability following genotoxic stress.

The spatial regulation of EXD2 activity also highlights an important principle in genome maintenance pathways. Nucleases capable of processing nucleic acid intermediates are often tightly controlled through subcellular localization, post-translational modifications, or regulated recruitment to specific substrates (*51*), (*52*), (*53*). In the case of EXD2, mitochondrial sequestration under basal conditions may prevent inappropriate nuclear nuclease activity. Consistent with this idea, persistent nuclear localization of catalytically active EXD2 leads to chromosome alignment defects and micronucleus formation, indicating that uncontrolled nuclear nuclease activity can itself compromise genome integrity.

Several cellular pathways contribute to the resolution of RNA–DNA hybrids, including RNaseH1/H2 enzymes, helicases such as senataxin, and multiple RNA processing factors (*50*). The requirement for EXD2 in this context suggests that it fulfills a specialized role in resolving R-loop–like intermediates generated at elongation-arrested RNAPII complexes during transcription shutdown. Whereas RNaseH enzymes cleave RNA within R-loops, EXD2 may instead trim RNA molecules associated with stalled transcription complexes, potentially facilitating their subsequent removal by canonical hybrid-processing pathways. This model would explain why EXD2 activity becomes particularly important under conditions of genotoxic stress that promote widespread transcriptional arrest.

Finally, our findings raise the possibility that stress-induced nuclear trafficking pathways may represent a therapeutic vulnerability. Pharmacological inhibition of nuclear import enhances the cytotoxicity of the transcription-blocking chemotherapeutic agent cisplatin in cell-based assays. These observations suggest that limiting nuclear deployment of EXD2 may exacerbate transcription-associated genome instability in damaged cells. Beyond establishing EXD2 nuclear import as a general stress response, our results point to the potential of targeting nuclear trafficking pathways to sensitize cancer cells to transcription-blocking DNA-damaging agents.

In summary, our study reveals that stress-induced nuclear import of EXD2 coordinates the resolution of transcription-associated R-loop–like intermediates with the restoration of transcription following DNA damage. By linking spatial regulation of nuclease activity to transcription restart and genome stability, these findings uncover an unexpected mechanism through which cells coordinate transcription and DNA repair during the DNA damage response.

## Material and methods

### Cell lines

U-2 OS, U-2 OS GFP::EXD2 (WT) and GFP::EXD2 (mutNLS) (clones 1 and 2) cells were cultured in Dulbecco’s Modified Eagle Medium (DMEM) containing 1 g L^-1^ glucose, supplemented with 10% fetal calf serum (FCS) and gentamicin (40 μg mL^-1^). Cells were maintained at 37°C in a humidified atmosphere with 5% CO_2_. HeLa cell lines, including HeLa EXD2(+/+), EXD2(-/-), EXD2(-/-) + EXD2(ΔMTS) and EXD2(-/-) + EXD2(ΔMTS/ND), were cultured under the same conditions. For HeLa EXD2(-/-) + EXD2(ΔMTS) and EXD2(-/-) + EXD2(ΔMTS/ND) cells the medium was additionally supplemented with 0.25μg mL^-1^ of puromycin to maintain selection.

### CRISPR/Cas9

U-2 OS EXD2(WT)::GFP cells expressing EXD2(WT)::GFP from the endogenous EXD2 locus were prepared by genome editing using the CRISPR-Cas9 system, as follows. A guide RNA targeting DNA cleavage next to the STOP codon of the *EXD2* gene was ordered from Integrated DNA Technologies (targeted sequence, with guide RNA spacer sequence in uppercase and PAM in lowercase 5′AAAGCAGCTATCAAGA CAGCtgg3′). A donor plasmid was constructed with eGFP coding sequences inserted at the position of the *EXD2* STOP codon and flanked by the 5′ homology arm chr14:69 240 298-69 241 097 (hg38) and 3′ homology arm chr14:69 241 101-69 241 897 (hg38). Guide RNA and Cas9 protein were assembled into RNP, mixed with donor plasmid and electroporated into U-2 OS cells using the Amaxa nucleofector II with kit V and program X-001 (Amaxa, Lonza, Switzerland). Electroporated cells were treated for 48 h with 5 μM AZD-7648 to inhibit NHEJ and favor HDR-mediated repair leading to targeted GFP cDNA insertion. Clones of cells expressing EXD2(WT)::GFP were isolated after FACS sorting for GFP-positive cells and targeted insertion was confirmed by PCR amplification of the edited locus with 5’CCATTCACACAGCATCGGAG3’ and 5’TGCAACCTGACCAGATGACT3’ (corresponding to sequences outside the homology arms). A clone expressing EXD2(WT)::GFP and no longer expressing EXD2(WT) was used for further experiments. U-2 OS EXD2(mutNLS)::GFP cells expressing EXD2(mutNLS)::GFP were prepared from the U-2 OS EXD2(WT)::GFP cells described above. A guide RNA targeting DNA cleavage next to the candidate NLS sequence of the *EXD2* gene and single stranded donor DNA were ordered from Integrated DNA Technologies (guide RNA target sequence: 5′ AACATAAAAGAAAGCCTCTGggg 3′, with spacer sequence in uppercase and PAM in lowercase; donor DNA sequence: AGTCTAAGATGGATGGGATGGTGCCAGGCAACCACCAAGGGAGAGACCCCGCCGCTGCCGCAGCCGCTGCACTGGGGGTGGGCTATTCTGCCAGGTAACTGAATCACTCCCTGTCTAGTAA). Guide RNA and Cas9 protein were assembled into RNP, mixed with donor DNA and electroporated into U-2 OS EXD2(WT)::GFP cells as above. Three days after electroporation, cells were sorted and seeded at 1 cell/well in a 96 well plate. Clones were analyzed by PCR with primers TCAGTGGCTCTCTTTCTTCATCT and AAGCAAGATTAACCCCGGGA and Sanger sequencing. Two independent clones with expected *EXD2(*mutNLS) sequence and no longer having *EXD2(WT)* sequence at the candidate NLS were used for further study.

### Transfection of cells

siRNA transfections were performed using Lipofectamine RNAiMAX Reagent (Invitrogen), according to the manufacturer’s instructions. Plasmid transfections (wtRNaseH1-GFP, dRNaseH1-GFP and Homologous recombination plasmids) were performed using X-tremeGENE9 DNA Transfection Reagent (Roche), according to the manufacturer’s instructions.

### Cell survival

Cells were grown on coverslips in 6-well plates and treated with the indicated treatments, including drug exposure and transfection according to the experimental design. Cells were exposed to UV irradiation using a UV lamp at doses ranging from 0 to 30 J m^-2^. After UV irradiation, cells were incubated for 48 h at 37 °C in a humidified atmosphere with 5% CO_2_, in the continued presence of the respective treatments. After the recovery period, cells were fixed with 4% paraformaldehyde (PFA) for 10 min at room temperature (RT) and stained with 0.5% crystal violet solution for 20 min at RT under agitation. Cells were washed, air-dried at RT overnight and then incubated with 1% SDS under agitation at RT until complete solubilization of the dye. Absorbance was measured at 590 nm to quantify cell viability.

### EU staining

Cells were grown on coverslips in 24-well plates and treated with the indicated treatments, including drug exposure, UV-irradiation (20 J m^-2^) and/or transfection according to the experimental design. Following drug treatments and/or UV irradiation, cells were incubated with 0.1 mM 5-ethynyl uridine (5EU) for either 1 h (for RRS experiments) or 10 min (for RNA degradation assays). Cells were then fixed with 4% PFA for 10 min at RT. Additionally, cells were treated with 0.05 µg mL^-1^ actinomycin D, added 30 min prior to and maintained during 5EU incorporation to inhibit RNAPI transcription. Subsequent steps were carried out using the Click-iT™ RNA Alexa Fluor™ 488 or 594 Imaging Kit (Invitrogen, C10329) according to the manufacturer’s protocol. Nuclei were counterstained with freshly prepared DAPI (1 μg mL^-1^ in PBS) for 2 min at RT. Cells were mounted with ProLong™ Gold antifade reagent (Molecular Probes). Fluorescence images were acquired using a Leica DM 6 B microscope equipped with a Hamamatsu OCRA-Flash4.0LT camera. Quantification of 5EU signal intensity was performed using Fiji (ImageJ) software.

### Immunofluorescence-based DNA lesions quantification

Cells were grown on coverslips in 24-well plates and exposed or not to UV irradiation (20 J m^-2^), followed by recovery times from 0 to 6 h. Cells were then fixed with 4% PFA for 10 min at RT and permeabilized with 0.5% Triton X-100 for 15 min at RT under gentle agitation. DNA denaturation was then performed with 2 M HCl for 20 min at RT and cells were blocked in 10% heat-inactivated fetal calf serum (FCS) in PBS for 1 h, at RT under agitation. Immunostaining of DNA photoproducts was performed using mouse monoclonal anti-6-4PP antibody diluted in 10% FCS in PBS and incubated for 2 h, at RT. After three washes in 0.1% Triton X-100 in PBS (5 min each under agitation), cells were incubated with the secondary antibody diluted in 10% FCS in PBS, for 1 h, at RT. Final washes were carried out in 0.1% Triton X-100 in PBS (3 x 5 min under agitation). Nuclei were counterstained with freshly prepared 1 μg mL^-1^ DAPI in PBS for 2 min at RT and cells were mounted using the ProLong™ Gold antifade reagent (Molecular Probes). Fluorescence images were acquired using a Leica DM 6 B microscope equipped with a Hamamatsu OCRA-Flash4.0LT camera. In each experiment, images of the cells were obtained with the same microscopy system and constant acquisition parameters. Signal intensities were quantified using Fiji software (ImageJ) to assess the percentage of 6–4PP removal (100% corresponding to the level of lesions detected immediately after UV irradiation).

### Immunofluorescence

Cells were grown on coverslips in 24-well plates and treated with the indicated treatments, including drug exposure and UV-irradiation (20 J m^-2^) according to the experimental design. Cells were fixed with 4% PFA for 10 min at RT. For EXD2 localization experiments, cells were incubated with MitoTracker (250 nM) for 30 min prior to fixation to stain mitochondria. Following fixation, cells were permeabilized with 0.1% Triton X-100 for 3 x 5 min at RT under gentle agitation. Blocking was performed in 10% heat-inactivated FCS in PBS for 30 min at RT under agitation. Primary antibodies were carried out for 1 h, at RT in a solution containing 10% heat-inactivated FCS and 0.1% Triton X-100 in PBS, followed by washes with PBS 0.1% Triton X-100 for 3 × 5 min at RT under agitation. Secondary antibodies, diluted in 10% FCS in PBS, were applied for 1 h, at RT, followed by washes in 0.1% Triton X-100 in PBS (3 x 5 min under agitation). Nuclei were counterstained with freshly prepared 1 μg mL^-1^ DAPI in PBS for 2 min at RT and cells were mounted using ProLong™ Gold antifade reagent (Molecular Probes). Fluorescence images were acquired using a Leica DM 6 B microscope equipped with a Hamamatsu OCRA-Flash4.0LT camera and quantification of EXD2 signal intensity was performed using Fiji software (ImageJ). Representative images were taken with inverted Leica SP8 microscope. In each experiment, images of the cells were obtained with the same microscopy system and constant acquisition parameters.

### Time course quantification of nuclear EXD2

Cells were grown on coverslips and UV-irradiated at 20 J m^-2^. After irradiation, cells were incubated at 37°C with 5% CO_2_ for different periods of time, washed with PBS at RT, and fixed with 2% PFA for 15 min at 37° C. Cells were permeabilized with PBS 0.1 % Triton X-100 (3X short + 2x 10 min washes). Blocking of non-specific signals was performed with PBS+ (PBS, 0.5 % BSA, 0.15 % glycine) for at least 30 min. Then, coverslips were incubated with a primary antibody against GFP diluted in PBS+ for 2 h, at RT in a moist chamber. After several washes with PBS + 0.1% Triton X-100 (3x short + 2x 10 min) and a quick wash with PBS+, cells were incubated for 1 h, at RT in a moist chamber with the secondary antibody coupled to fluorochrome and diluted in PBS+. After the same washing procedure but with PBS, coverslips were finally mounted using Vectashield with DAPI (Vector Laboratories). Confocal images of the cells were obtained on a Plan-Apochromat 40x/1.3 oil immersion objective. Images of the cells for each experiment were obtained with the same microscopy system and constant acquisition parameters. Images were analyzed with Fiji software (ImageJ).

### Mitotic defects and micronuclei quantification

Mitotic defects: cells were grown on coverslips in 24-well plates and treated with 15 μM RO-3306, an inhibitor of CDK1, for 18 h to induce G2/M arrest, followed by drug release for 1 h to allow cell cycle progression. After the release period, cells were fixed with 4% PFA for 10 min at RT. Immunofluorescence staining was then performed using an anti-α-tubulin antibody, following the standard protocol described above. Mitotic defects were quantified by blind counting of at least 300 cells per condition.

Micronuclei: cells were grown on coverslips in 24-well plates and fixed with 4% PFA for 10 min at RT. Nuclei were counterstained with 1 μg mL^-1^ DAPI in PBS for 2 min at RT and cells were mounted using ProLong™ Gold antifade reagent (Molecular Probes). Confocal microscopy images were acquired in a z-stack mode with a Leica SP8 microscope. Micronuclei analysis has been made with Fiji software (ImageJ) and for each field, the number of micronuclei was divided by the number of nuclei.

### Proximity ligation assay

Cells were grown on coverslips in 24-well plates and treated with the indicated treatments, including drug exposure, UV-irradiation (20 J m^-2^) and/or transfection according to the experimental design. Cells were fixed with 4% PFA for 10 min at RT, followed by a permeabilization in 0.1% Triton X-100 in PBS for 15 min at RT under gentle agitation. The PLA was then performed using the Duolink® Proximity Ligation Assay kit (Sigma-Aldrich) according to the manufacturer’s protocol. Antibody incubations were performed for 1 h, at RT. The following antibody pairs and host species were used: RNAPII (mouse) / EXD2 (rabbit), Ser5P/Ser2P (mouse) / EXD2 (rabbit), RNAPII (mouse) / GFP (rabbit) and flag (mouse) / GFP (rabbit). Antibody dilutions are provided in the Key resources table. For negative controls, each experiment included samples incubated with only one of the antibodies. Nuclei were counterstained with freshly prepared DAPI (1 μg mL^-1^ in PBS) for 2 min at RT and cells were mounted using the ProLong™ Gold antifade reagent (Molecular Probes). Fluorescence images were acquired using a Leica DM 6 B equipped with a Hamamatsu OCRA-Flash4.0LT camera. Image analysis and quantification of PLA signals per nucleus were performed using Fiji software (ImageJ).

### Homologous Recombination assay

Cells were grown on coverslips in 6-well plates and treated with IVM (1 μM) or DMSO for 1 h. Homologous Recombination (HR) repair activity was assessed using the Homologous Recombination Assay kit (Norgen Biotek Corporation) according to the manufacturer’s protocol. Cells were transfected with the plasmid-based HR reporter system provided in the kit using XtremGen9 transfection reagent. After 20 h recovery period, genomic DNA was extracted using the Monarch Spin gDNA Extraction kit (New England Biolabs). HR-mediated recombination events were quantified by quantitative PCR (qPCR) using the primer sets supplied with the kit and SYBR Green Master Mix on a LightCycler 96 instrument (Roche). qPCR amplification of the recombinant product was used to determine HR efficiency according to the manufacturer’s guidelines.

### Whole cell extract preparation and western blot

Cells were harvested and resuspended in RIPA buffer (10 mM Tris-HCl, pH 8; 1 mM EDTA; 1 mM EGTA; 1% triton X-100; 0.1% sodium deoxycholate; 0.1% SDS; 140 mM NaCl). After a 30 min incubation at 4°C under agitation, a centrifugation was performed at 16,000 x g for 20 min at 4°C.

### Synergy assay

Cells were seeded in 96-well plates and were treated for 72 h with increasing concentrations of CisPt (0,01 – 0,5 µM) and IVM (0,5 – 20 µM) either as single agents or in combination. PrestoBlue Cell Viability Reagent (Thermo Fisher Scientific) was used to quantify cell viability according to the manufacturer’s instructions. The absorbance was measured with a CellInsight CX5 High Content Screening platform (Thermo Fisher Scientific) at 600 nm, and viability was expressed as percentage relative to DMSO-treated controls. Dose–response curves were generated from cell viability assays. Synergy Scores were calculated using the Highest Single Agent (HSA) model via the SynergyFinder+ web tool (*54*). According to the HSA model, drug combinations with synergy scores < 0 are considered antagonistic, = 0 additive, and > 0 synergistic, with higher positive or negative values indicating stronger synergy or antagonism, respectively.

### Statistics and reproducibility

Experimental data were plotted and analyzed using GraphPad Prism (GraphPad Software Inc.). The number of samples and replicates are indicated in the respective figure legends. Each experiment was repeated at least twice with similar results. When indicated Standard Error of the Mean (SEM) is shown.

## Funding

This study was supported by the Ligue contre le cancer (Equipe Labélisée 2022-2024), the Institut National du Cancer (INCa) (INCa_18353), the grant ANR-20-CE12-0017 (TFIIH), ANR-22-CE12-0019 (MitoRare), ANR-24-CE12-1416 (BAF-NER), ANR-24-CE12-7268 (Nu-DyRection), The “Fondation pour la Recherche Médicale) (FRM EQU202603051215), ANR-10-LABX-0030-INRT, a French State fund managed by the Agence Nationale de la Recherche under the frame program Investissements d’Avenir ANR-10-IDEX-0002-02. This work of the Interdisciplinary Thematic Institute IMCBio+, as part of the ITI 2021-2028 program of the University of Strasbourg, CNRS and Inserm, was supported by IdEx Unistra (ANR-10-IDEX-0002), and by SFRI-STRAT’US project (ANR-20-SFRI-0012) and EUR IMCBio (ANR-17-EURE-0023) under the framework of the France 2030 Program. This work was supported by the Fondation pour la Recherche Médicale, grant number «ECO202406019123 », to CC. We acknowledge the IGBMC imaging center, member of the national infrastructure France-BioImaging supported by the French National Research Agency (ANR-10-INBS-04). We also thank the IGBMC PluriCellEast and Photonic microscopy services.

## Author contributions

Conceptualization: JS and FC

Methodology: JS and FC

Investigations: JS, PC, CC, AH, EB, LMD, POM, JPC and AB

Analysis of data: JS and FC

Funding acquisition: FC, EB

Supervision: FC

Writing – original draft: FC

Writing – review & editing: FC, JPC, EB and JS

## Data and material availability

Further information and requests for resources and reagents should be directed to and will be fulfilled by Frédéric Coin.

Sup. Table 1 contains a full list of antibodies, oligonucleotides, cell lines and reagents used in this study with RRID reference numbers when available.

## Sup. Fig. Legends

**Sup. Fig. 1.**
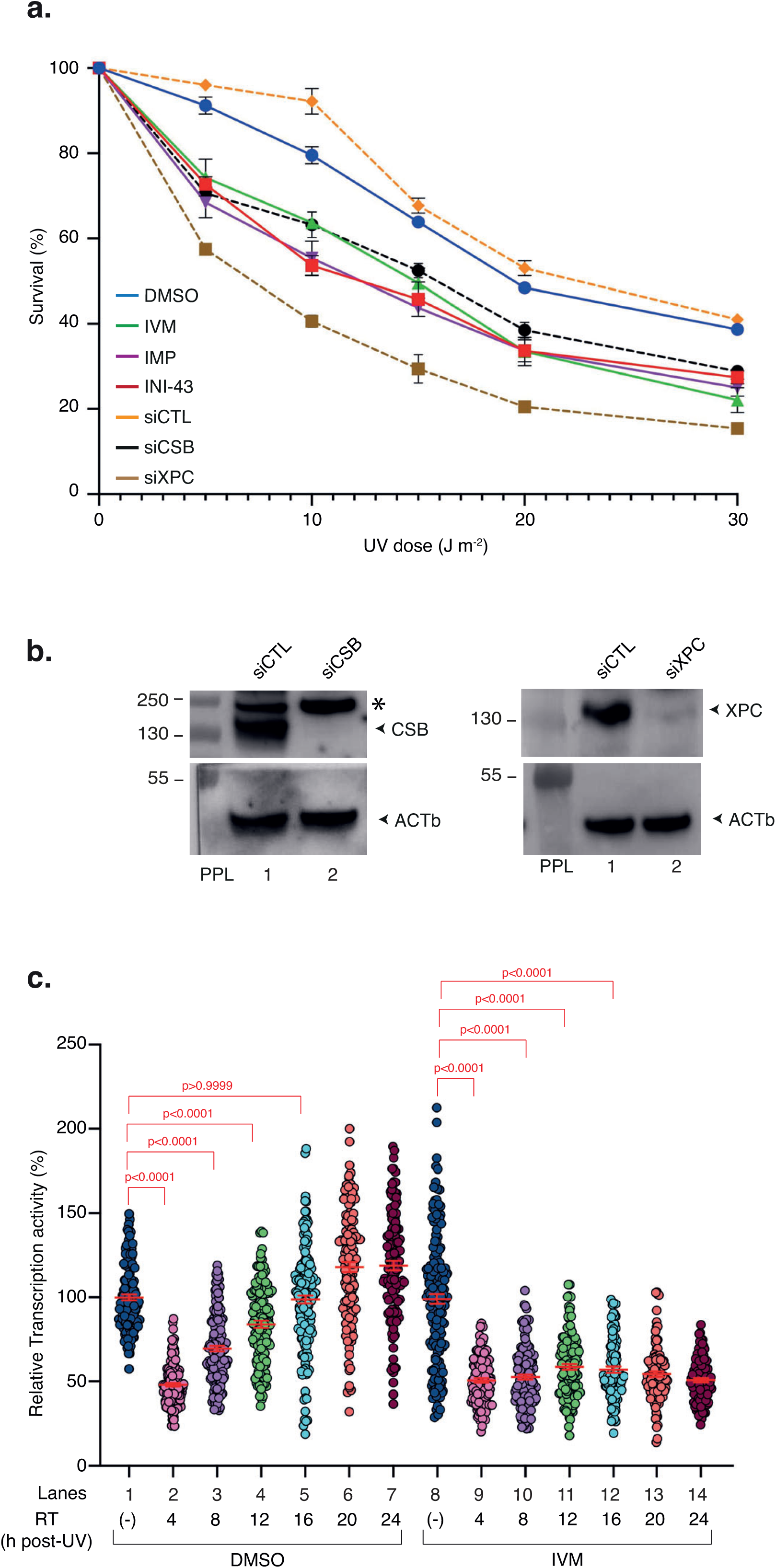
Importin-α/β1-dependent nuclear import sensitizes cells to UV irradiation (a), Cells were treated with increasing doses of UV irradiation and survival was determined 48 h later. Data were normalized to the mock treatment controls (set at 100%). Data are mean ± s.e.m. from three independent experiments. When indicated, before UV irradiation, cells were treated for 1 h with inhibitors of nuclear import (1μM) or for 48 h with siCTL, siCSB or siXPC. **(b),** Protein lysates of U-2 OS cells treated with siCTL, siXPC or siCSB for 48 h were immuno-blotted for XPC, CSB or ACTb as indicated. Molecular weights (kDa) and prestained protein ladders (PPL) are shown. (*), non-specific band. **(c),** U-2 OS cells were mock-or UV-irradiated (20 J m^-2^) and mRNA was labeled with EU at the indicated time points post-UV. When indicated, cells were treated for 1 h with IVM (1μM) before UV irradiation. To ensure specificity for RNAPII, cells were pre-treated with a low dose of actinomycin D to inhibit RNAPI-driven transcription. EU signal was quantified by ImageJ and relative integrated densities, normalized to mock-irradiated level set to 100%, are reported on the graph (*n*>300 cells per condition from three independent experiments). Red bars indicate mean integrated densities ± s.e.m.. RT; recovery time. (-); cells were mock-irradiated. Statistical analyses were performed using one-way ANOVA with Tukey’s post hoc test.

**Sup. Fig. 2.**
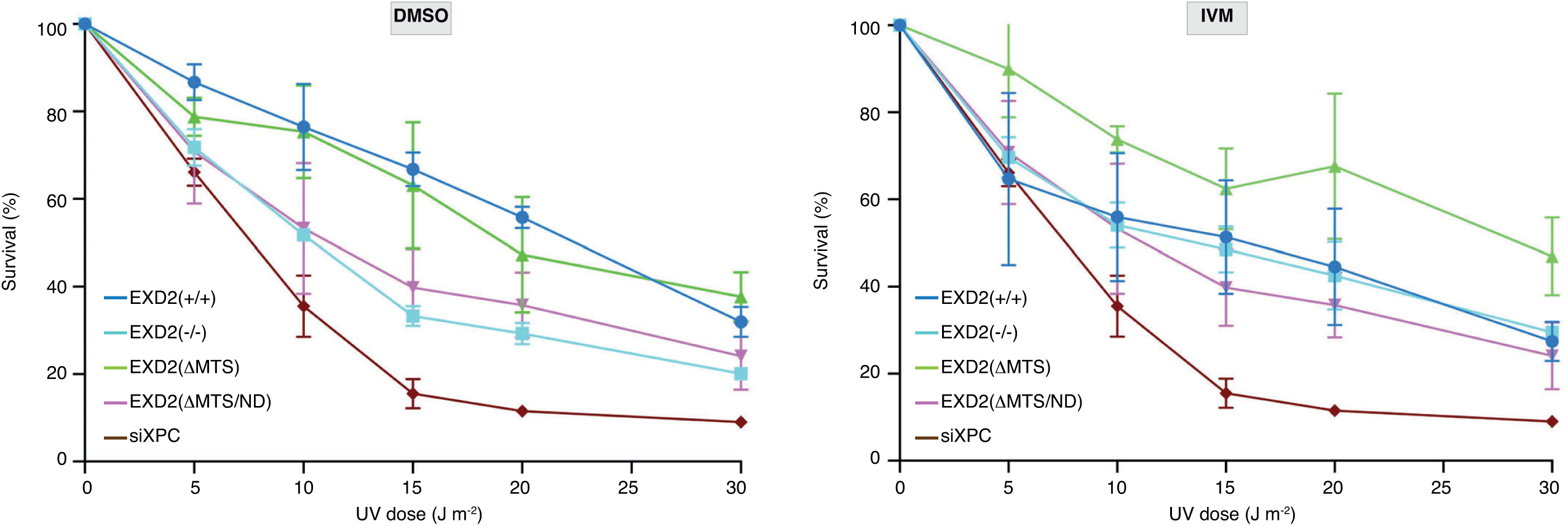
Forced EXD2 nuclear localization circumvents nuclear import inhibition of UV survival. HeLa EXD2(+/+), EXD2(-/-), EXD2(ΔMTS) and EXD2(ΔMTS/ND) cells were treated with increasing doses of UV irradiation and survival was determined 48 h later. Cells were treated either with DMSO (left) or IVM (1 μM) (right). When indicated, before UV irradiation, cells were treated for 24 h with siXPC. Data were normalized to the mock-irradiated controls (set to 100%). Data are mean ± s.e.m. from three independent experiments. Statistical analyses were performed using one-way ANOVA with Tukey’s post hoc test.

**Sup. Fig. 3.**
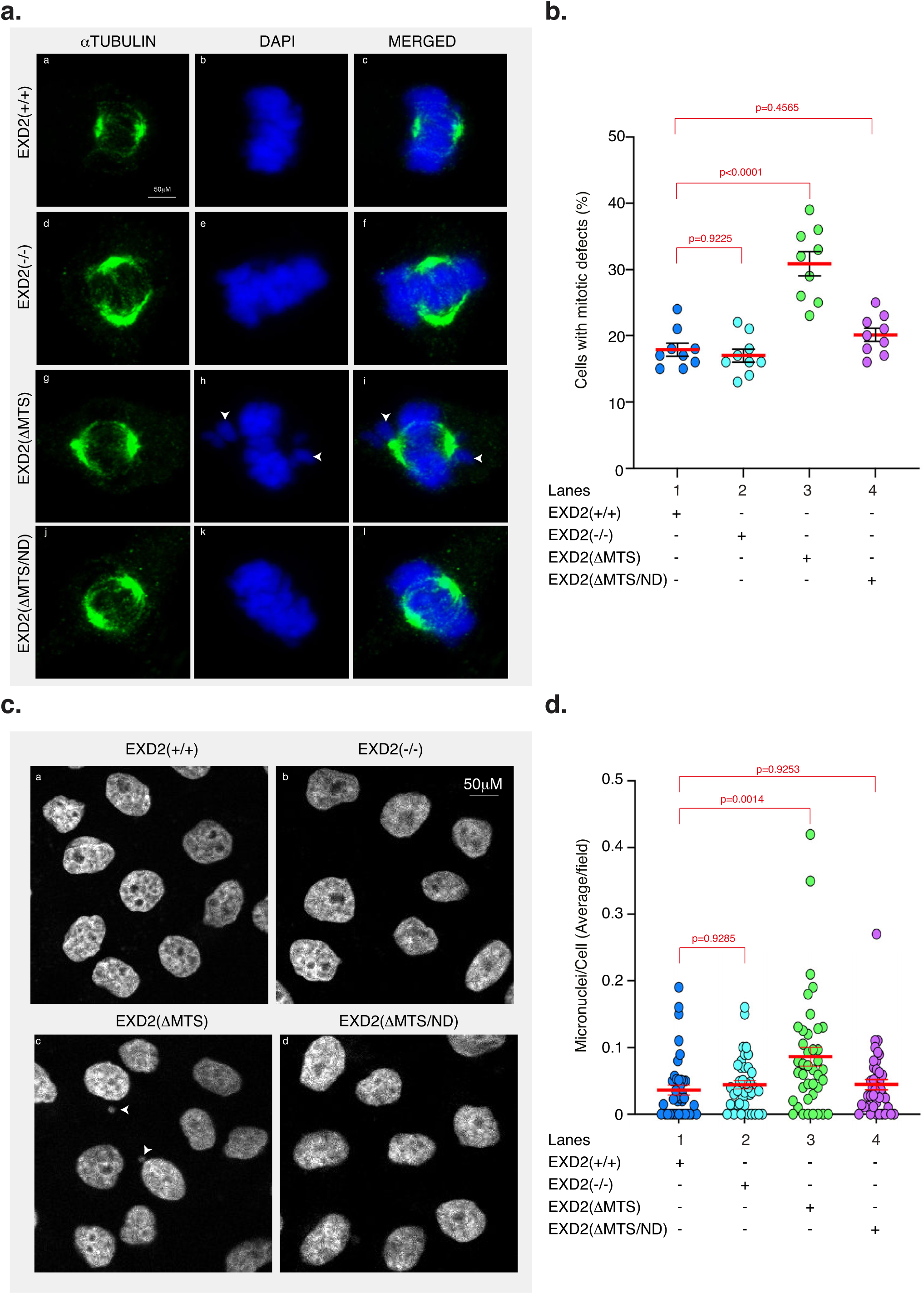
Persistent nuclear localization of EXD2 causes chromosome misalignment and micronuclei formation (a), Representative confocal images of HeLa EXD2(+/+), EXD2(-/-), EXD2(ΔMTS) and EXD2(ΔMTS/ND) cells analyzed in mitosis. Cells were treated with RO-3306, an inhibitor of CDK1, for 18 h. After a release for 60 min, cells were fixed and labeled with anti-αTubulin. The arrows point to chromosome segregation errors at metaphase in HeLa EXD2(ΔMTS) cells. MERGED: overlay of αTubulin labeling and DAPI staining. **(b),** Quantification of mitotic cells displaying either a normal or a defective mitotic phenotype (*n* > 300 cells per condition from three independent experiments). **(c),** Representative confocal images of HeLa EXD2(+/+), EXD2(-/-), EXD2(ΔMTS) and EXD2(ΔMTS/ND) cells analyzed for the presence of micronuclei. The arrows point to micronuclei in HeLa EXD2(ΔMTS) cells. **(d),** Quantification of micronuclei per cell in HeLa EXD2(+/+), EXD2(-/-), EXD2(ΔMTS) and EXD2(ΔMTS/ND) cells (*n* = 40 fields per condition from three independent experiments). Statistical analyses were performed using one-way ANOVA with Tukey’s post hoc test.

**Sup. Fig. 4.**
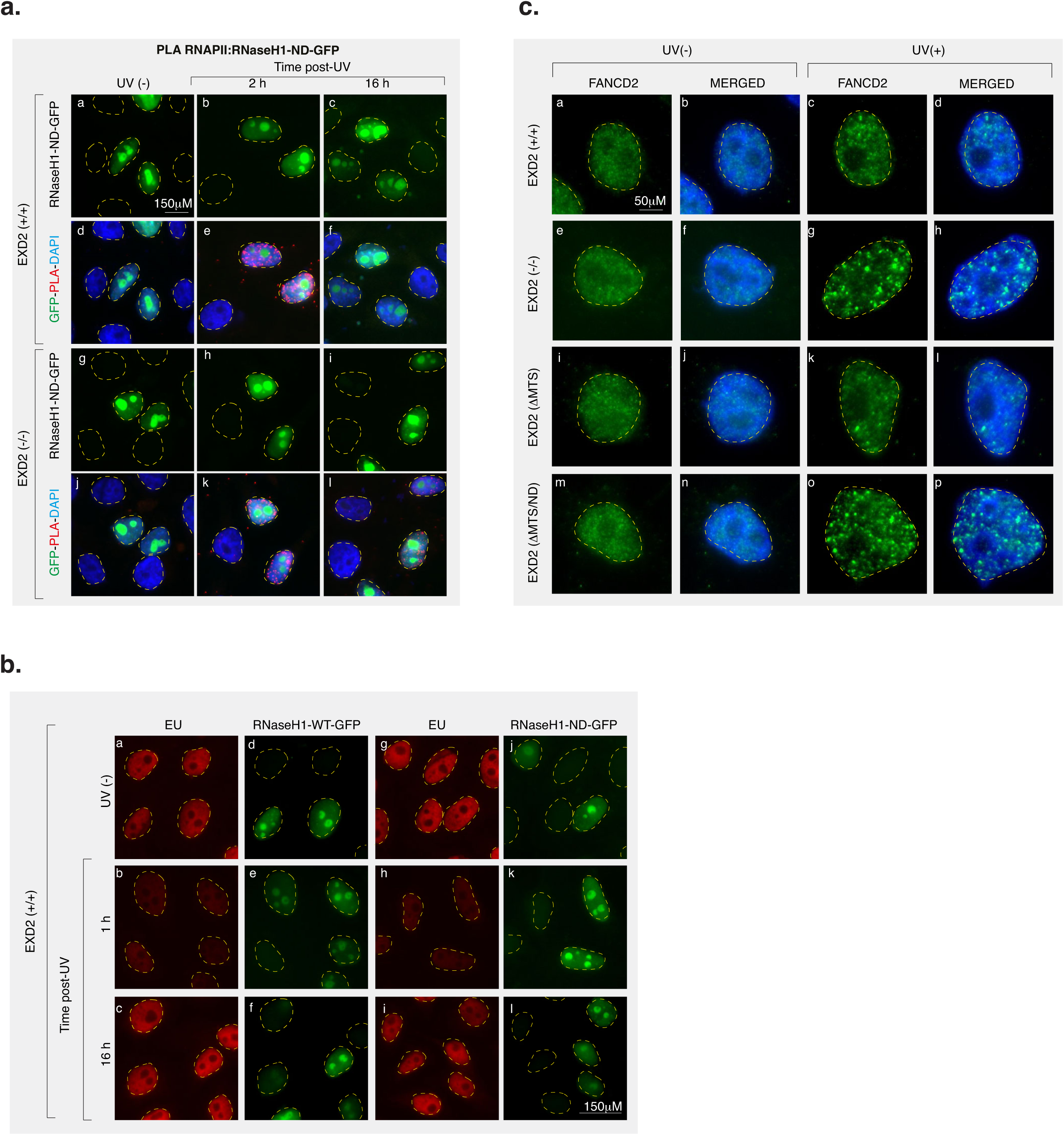
EXD2 is involved in the resolution of R-loop intermediates associated with RNAPII complexes after UV irradiation (a), Representative microscopy images of PLA between RNAPII and dRNaseH1-GFP performed in HeLa EXD2(+/+) and EXD2(-/-) cells mock-or UV-irradiated (20 J m^-2^) and let to recover for 2 or 16 h. **(b),** Representative microscopy images of EU incorporation assay in HeLa EXD2(+/+) cells expressing either wtRNaseH1-GFP or an inactive dRNaseH1-GFP (green) mock-or UV-irradiated at 20 J m^-2^.Nascent RNA synthesis was monitored by EU incorporation (red) at the indicated times following UV irradiation. **(c),** Representative microscopy images of FANCD2 foci (green) in HeLa EXD2(+/+), EXD2(-/-), EXD2(ΔMTS) and EXD2(ΔMTS/ND) cells mock-or UV-irradiated at 20 J m-^2^ (recovery time; 16h). After recovery time, cells were fixed and labeled with anti-FANCD2. Nuclei were counterstained with DAPI (blue). MERGED: overlay of FANCD2 labeling and DAPI staining.

**Sup. Fig. 5.**
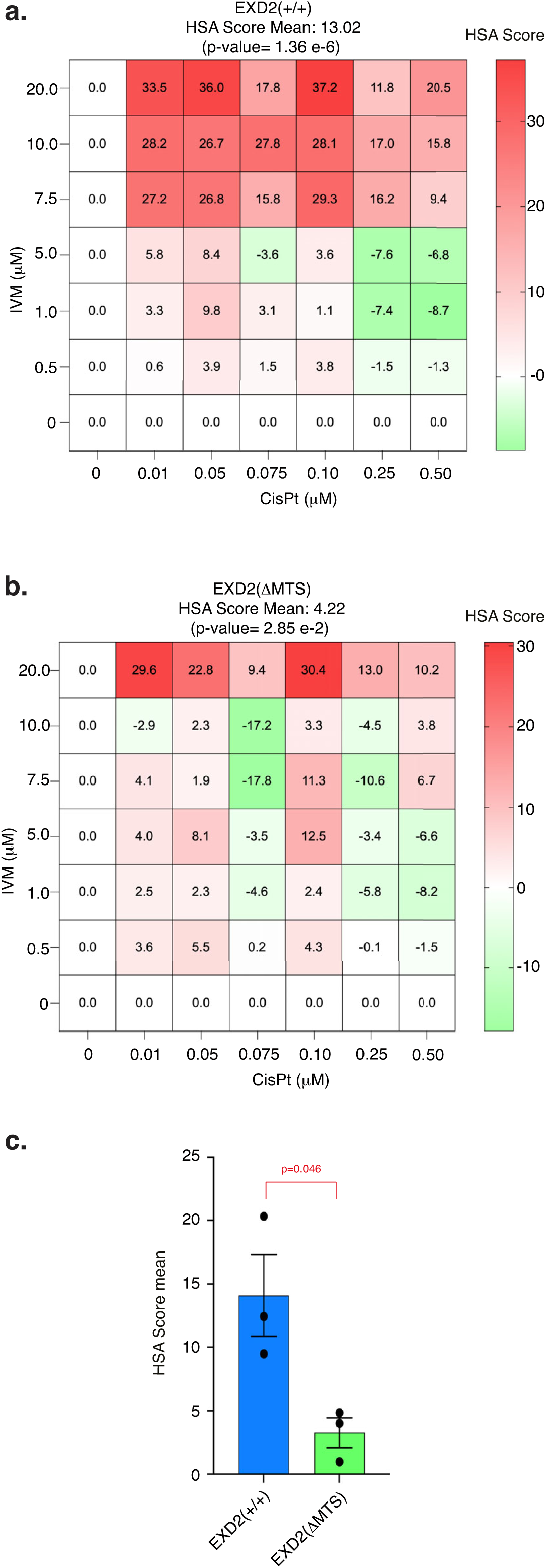
Pharmacological inhibition of EXD2 nuclear import enhances the efficacy of transcription-blocking chemotherapeutic agent CisPt **(a,b),** Drug–drug interaction analysis of IVM and CisPt in HeLa EXD2(+/+) cells **(a)** or HeLa EXD2(ΔMTS) **(b)**. Heatmaps show Highest Single Agent (HSA) synergy scores across the indicated drug concentrations. Mean HSA scores and associated p values are shown above each heatmap. **(c),** Quantification of mean HSA scores for IVM + CisPt combinations in EXD2(+/+) and EXD2(ΔMTS)-expressing cells. Data are mean ± s.e.m. from three independent experiments. Statistical significance was assessed using an unpaired two-tailed t-test.

---

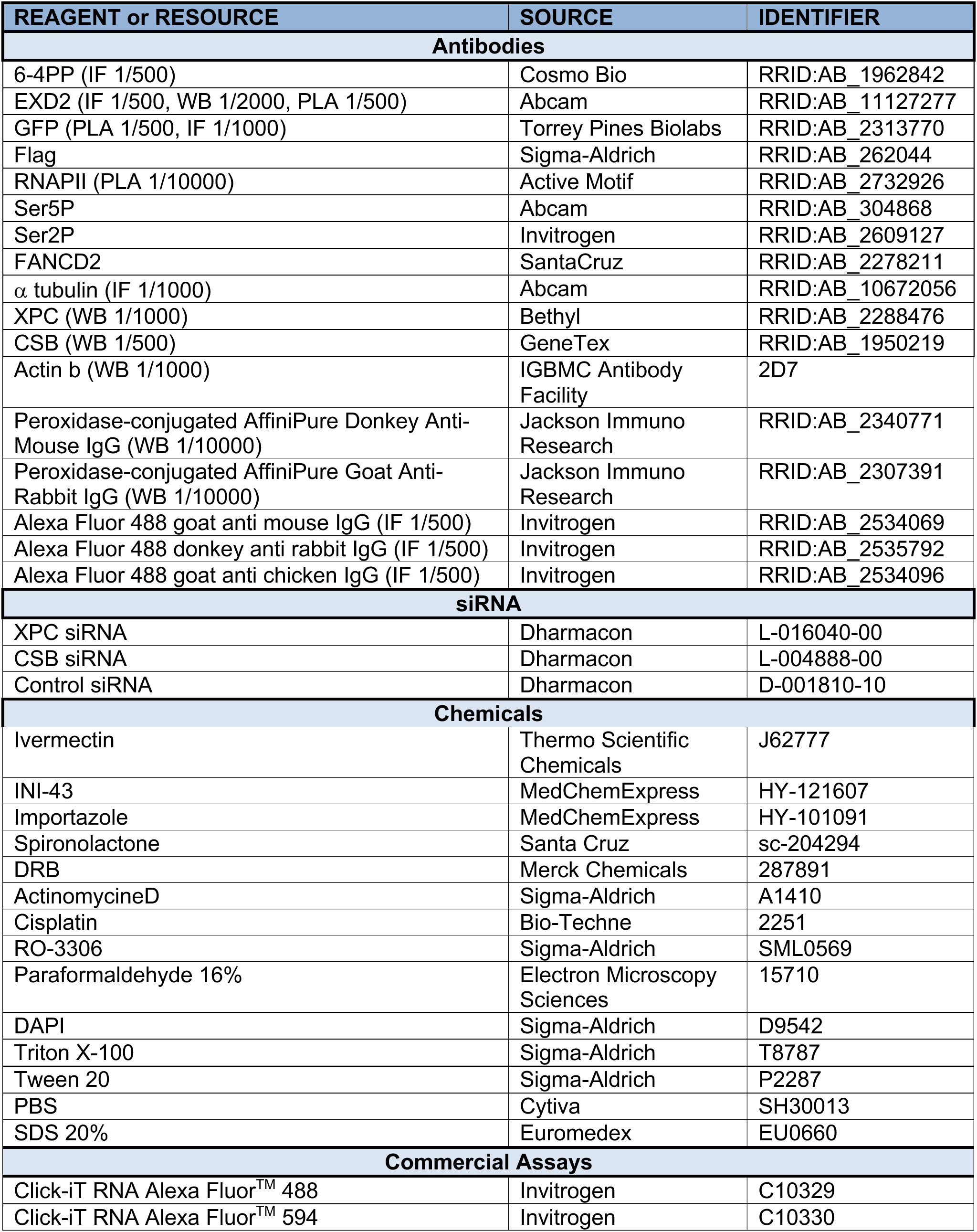
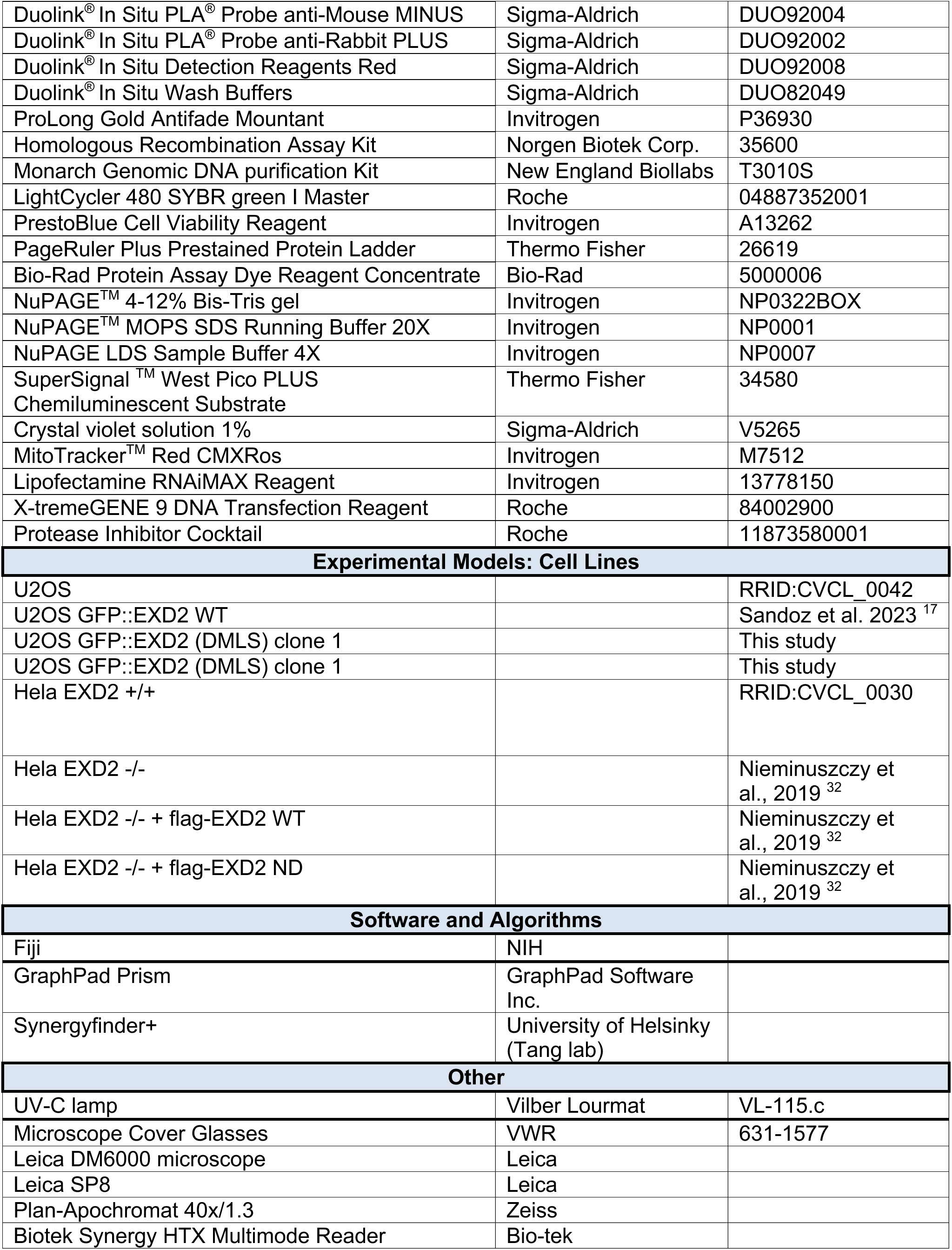
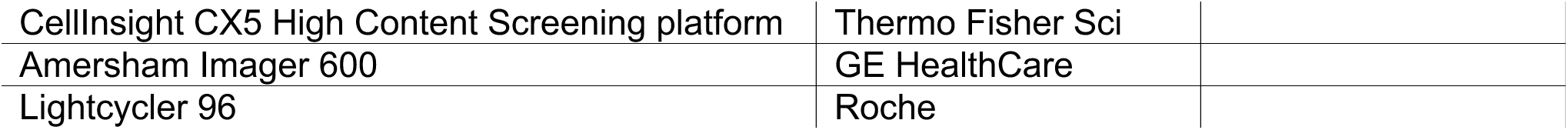

## Notes

https://www.researchsquare.com/article/rs-7241921/v1

### Competing Interest Statement

The authors have declared no competing interest.

### Summary of Updates

several experiments have been added to strengthen the message the discussion has been modified accordingly

